# Intersectional genetic targeting of molecularly distinct melanin-concentrating hormone neuron subpopulations reveals distinct and overlapping subcircuit architecture

**DOI:** 10.64898/2026.10.05.756884

**Authors:** Monica S. Antony, Persephone A. Miller, William P. Armstrong, Leo DiLeone, Olivia Knuth, Mishael Gill, Tammy Luong, Elizabeth Chamiec-Case, Arnab Barik, Alexander T. Chesler, Lindsay A. Schwarz, Melissa J. Chee, Alexander C. Jackson

## Abstract

The lateral hypothalamic area (LHA) coordinates crucial and innate behaviors through heterogeneous yet poorly understood neuronal populations and circuits. Neurons that produce melanin-concentrating hormone (MCH; encoded by the *Pmch* gene) are primarily localized in the LHA and contribute to feeding, sleep-wake regulation, memory, and reward-associated behaviors. Building on foundational anatomical studies, our single-cell transcriptomic analysis of the LHA identified markers distinguishing at least two transcriptionally distinct MCH subpopulations. These subpopulations are distinguished by the selective enrichment of the *Tacr3* and *Cartpt* transcripts, with each comprising approximately half of the total MCH population. However, these subpopulations remain poorly understood owing to limited genetic targeting tools. We hypothesized that MCH neurons operate through functionally distinct, parallel subcircuits that exhibit unique neuroanatomical projections. In this study, we leveraged *Pmch* and *Tacr3* expression to intersectionally dissociate these neurons into Type I MCH (*Pmch+/Tacr3+*) and Type II MCH (*Pmch+/Tacr3-*) subpopulations. We first validated a *Tacr3*-Flpo mouse line for the genetic targeting of the Type I MCH subpopulation. We then generated an intersectional Type I MCH neuron reporter line to visualize Type I axonal projections. In parallel, we injected a dual recombinase mouse line with the intersectional genetic targeting viruses, Conditional Viral Expression by Ribozyme Guided Degradation (ConVERGD), and identified overlapping and distinct projections by Type I and Type II MCH neurons using fluorescence microscopy, super-resolution imaging, and high-spatial resolution fiber mapping. Overall, these data establish the first subpopulation-resolved anatomical framework for molecularly distinct Type I and Type II MCH subcircuits, thus establishing a foundation for future circuit-level functional studies.

**KEY POINTS:** xxxxx

## INTRODUCTION

Melanin-concentrating hormone (MCH), encoded by the highly conserved *Pmch* gene, is primarily produced in neurons within the lateral hypothalamic area (LHA) (Bittencourt, 2011; Kawauchi et al., 1983). From the LHA, MCH neurons project broadly throughout the central nervous system to influence a diversity of innate physiological and behavioral processes (Bittencourt et al., 1992; Diniz & Bittencourt, 2017). Early studies of the MCH system established a role in energy balance, as intracerebroventricular (icv) administration of MCH increased feeding (Qu et al., 1996), and either MCH deletion (Shimada et al., 1998) or genetic ablation of MCH neurons (Alon & Friedman, 2006) resulted in lean mice, characterized by decreased metabolic activity and hyperactivity. Paradoxically, icv MCH administration increased rapid eye movement (REM) sleep (Verret et al., 2003), thus also implicating MCH in sleep regulation. Additionally, single-unit recordings identified MCH neurons as maximally active during REM sleep (Hassani et al., 2009), and *in vivo* optogenetic activation of MCH neurons extended REM sleep episodes (Jego et al., 2013; Konadhode et al., 2013; Tsunematsu et al., 2014). Today, we understand MCH neurons to be implicated in a broad variety of physiological behaviors, such as exploration (Blanco-Centurion et al., 2019; Concetti et al., 2024; González et al., 2016), reward-seeking (Dilsiz et al., 2020; Furman et al., 2025), memory consolidation (González et al., 2016; Izawa et al., 2019; Monzon et al., 1999), and non-homeostatic feeding (Furman et al., 2025; Subramanian et al., 2023).

1. The *Tacr3*-Flpo mouse line is specific in Flpo recombinase expression and can be used to generate a dual recombinase *Pmch*-Cre; *Tacr3*-Flpo mouse line for selective intersectional targeting of MCH subpopulations.
2. Type I and Type II MCH neurons exhibit both distinct and overlapping axonal projections with differential projections most evident in cortical and septal subregions.
3. MCH, CART, and VGLUT2 immunoreactivity are present in Type I and Type II MCH axons within the septal complex using super-resolution imaging.

One possible explanation for these diverse, yet contradictory, functions is that the LHA MCH population is comprised of functionally distinct subpopulations that are differentially recruited to mediate specific roles across behavioral states. Indeed, fiber photometry recording of MCH neuron calcium activity across sleep and wake states has revealed functionally distinct subpopulations: REM sleep-active, wake-active, and both REM and wake-active MCH neurons (Izawa et al., 2019). A parallel study also identified that a subset of REM-active MCH neurons is engaged during exploratory behaviors (Blanco-Centurion et al., 2019). Additionally, insulin receptor inactivation in MCH neurons leads to increased locomotor activity and improved insulin sensitivity in obese mice (Hausen et al., 2016) while inactivation of orexin receptor 2 in MCH neurons reveal shortened REM sleep with decreased insulin sensitivity in female mice (Izawa et al., 2025). These findings illustrate how MCH subpopulations may engage in different signaling modalities to influence downstream behavior, further supporting that these neurons are composed of functionally distinct subpopulations. We hypothesized that functionally distinct MCH subpopulations arise from molecularly distinct MCH subpopulations with both shared and unique axonal projection patterns, which may account in part for the diversity of sleep- and wake-related behaviors ascribed to the MCH system.

Early anatomical evidence revealed that MCH neurons are neurochemically heterogeneous and can be defined by the absence or presence of both cocaine- and amphetamine-regulated transcript (CART; encoded by *Cartpt*) (Broberger, 1999; Elias et al., 2001; Vrang et al., 1999) and neurokinin-3 receptor (NK3R; encoded by *Tacr3*) (Brischoux et al., 2002; Croizier et al., 2010; Cvetkovic et al., 2004; Griffond et al., 1997). Fate-mapping studies of these neurons using bromodeoxyuridine suggested that MCH subpopulations are developmentally distinct, with NK3R+/CART+ MCH neurons developing earlier than NK3R-/CART-MCH neurons (Brischoux et al., 2001, 2002). Retrograde True Blue labeling and immunostaining revealed that MCH subpopulations may also extend preferential anatomical projections, as NK3R+/CART+ MCH neurons strongly project towards the cortex, while NK3R-/CART-MCH neurons strongly project towards the brainstem and spinal cord (Croizier et al., 2010; Cvetkovic et al., 2004). These MCH subpopulations are intermixed within the LHA, but they exhibit some distinct spatial patterns. A greater proportion of NK3R+/CART+ MCH neurons are found in anterior and medial regions, and more NK3R-/CART-MCH neurons are found in posterior and lateral regions (Croizier et al., 2010; Miller et al., 2024; Wang et al., 2021). These subpopulations also exhibit distinct electrophysiological properties, with differences in firing rate and passive membrane properties (Adekunle et al., 2024; Fujita et al., 2021; Miller et al., 2024).

Building upon this evidence, our single-cell RNA sequencing of MCH neurons provided a transcriptional basis for the distinction between these subpopulations by revealing a suite of hundreds of differentially expressed genes (Mickelsen et al., 2019). We identified that *Tacr3* and *Cartpt* expression across MCH neurons is continuous rather than binary, yet the population can be divided into subpopulations based on the selective enrichment of these two genes. Overall, despite extensive developmental, anatomical, electrophysiological, and transcriptomic evidence supporting MCH heterogeneity, most functional studies manipulate MCH neurons as a single population, by necessity, owing to limited genetic tools available to selectively interrogate transcriptionally distinct MCH subpopulations.

To address this gap, we devised an intersectional genetic approach for interrogating molecularly distinct MCH subpopulations using two recombinase mouse lines targeting *Pmch* (*Pmch*-Cre) and *Tacr3* (*Tacr3*-Flpo). For the purposes of our study, we operationally defined Type I MCH neurons as expressing both markers (*Pmch+/ Tacr3+)* and Type II MCH neurons as expressing only *Pmch* (*Pmch+/Tacr3-*). This framework serves as a proxy for molecularly distinct MCH subpopulations. We first validated a novel knock-in *Tacr3*-Flpo mouse line to use *Tacr3* as a molecular toggle when accessing MCH subpopulations. Using this mouse line, we devised parallel but complementary intersectional strategies to assess Type I and Type II MCH neurocircuitry: a *Pmch*-Cre;*Tacr3*-Flpo;RC::FLTG mouse to label Type I MCH neurons, and a viral intersectional strategy using the *Pmch*-Cre;*Tacr3*-Flpo mouse line and utilizing ConVERGD (Conditional Viral Expression by Ribozyme Guided Degradation) (Hughes et al., 2024) viruses to label both Type I and Type II MCH axons. We then compared these subpopulation projections to total MCH projections, which we visualized using Cre-dependent anterograde viruses in the *Pmch*-Cre mouse. Using these strategies, we generated comprehensive maps of total, Type I, and Type II MCH fibers, with high resolution fiber mapping of MCH subpopulation fibers in the septal complex and adjacent cortex illustrating the divergent projection patterns by MCH subpopulations. Finally, using super-resolution microscopy, we defined MCH, CART and VGLUT2 immunoreactivity (IR) at virally labeled MCH subpopulation axons. Overall, these findings establish a subpopulation-resolved anatomical framework for genetically defined MCH circuitry and demonstrate that MCH neuronal subpopulations exhibit both overlapping and distinct projection patterns, providing a foundation for future investigations of MCH subcircuit function.

## MATERIALS AND METHODS

### Ethics statement

All experiments were performed under ethical guidelines established by the National Institutes of Health Guide for the Care and Use of Laboratory Animals and were approved by the Institutional Animal Care and Use Committee (protocol A25-055) at the University of Connecticut.

### Animals

Male and female mice were used for all experiments. All mice were group housed in a 12 hour light, 12 hour dark cycle and fed standard chow (Laboratory Rodent Diet 5001, LabDiet) and given water *ad libitum.* For manipulation of *Tacr3* neurons, we used the *Tacr3-*Flpo knock-in mouse line, generated using CRISPR-directed homologous recombination (Wu et al., 2018) to insert a T2A-FlpO cassette in frame with the 3’-end of *Tacr3*. For visualization of Flpo+ neurons, we crossed the heterozygous *Tacr3-*Flpo mouse line to homozygous Flpo-dependent Ai65 reporter line (B6;129S-Gt(ROSA)26Sortm65.1(CAG-tdTomato)Hze/J; The Jackson Laboratory (JAX) stock # 021875: RRID:IMSR_JAX:021875) (Daigle et al., 2018) to express tdTomato at *Tacr3* neurons in the *Tacr3*-Flpo;tdT mouse. For the manipulation of *Pmch* neurons, we used heterozygous *Pmch*-Cre (Tg(*Pmch*-cre)1Lowl/J transgenic mice; JAX stock #014099; RRID:IMSR_JAX:014099) (Kong et al., 2010). For intersectional experiments, we crossed the heterozygous *Pmch-*Cre mouse line to the heterozygous *Tacr3*-Flpo mouse line to generate a dual recombinase *Pmch-*Cre;*Tacr3*-Flpo animals. For intersectional fluorescent reporting of Type I MCH neurons, we crossed the dual recombinase *Pmch*-Cre;*Tacr3*-Flpo to the eGFP Cre/Flpo reporter line RC::FLTG (B6.Cg-Gt(ROSA)26Sortm1.3(CAG-tdTomato,-EGFP)Pjen/J; JAX stock # 026932: RRID:IMSR_JAX:026932)(Plummer et al., 2015) to express eGFP in *Pmch*- and *Tacr3*-positive neurons in the *Pmch*-Cre;*Tacr3*-Flpo; RC::FLTG mouse. The RC::FLTG breeders were donated by Dr. Natale Sciolino (Department of Physiology and Neurobiology, University of Connecticut, Storrs, CT).

### Stereotaxic surgery

For anterograde tracing of *Pmch* neurons, heterozygous *Pmch-*Cre animals were bilaterally injected (Nanoject III, Drummond Scientific) with 400 nl of AAV2-Ef1a-DIO-EYFP (Titer: 4.6 X 10E+12 GC/mL; University of North Carolina, Deisseroth Lab) into the LHA (in mm): anteroposterior (AP), −1.55; mediolateral (ML), ±.50, ±1.10; dorsoventral (DV), −4.90, −5.30. For Conditional Viral Expression by Ribozyme Guided Degradation (ConVERGD) (Hughes et al., 2024) mediated anterograde tracing of Type I and Type II MCH neurons, we injected either AAV2/8-hSyn-CV(Con/Fon)-eGFP-W3SL (Titer: 6.81E+12 GC/mL) or AAV2/8-hSyn-CV(Con/Foff)-eGFP-W3SL (Titer: 8.83E+12 GC/mL), respectively, into the LHA of *Pmch*-Cre;*Tacr3*-Flpo animals using the same coordinates and virus volume. ConVERGD viruses were gifted by Dr. Lindsay Schwarz (St. Jude Children’s Research Hospital).

### Transcardial perfusions, fixation and sectioning

Animals were anesthetized with vaporized isoflurane and a ketamine/xylazine/saline solution injected intraperitoneally. Animals were then transcardially perfused with 10 mL of .125 M saline, followed by 35 mL of 4% paraformaldehyde (PFA) in 1X PBS solution. Dissected brains were postfixed in 4% PFA in 4°C overnight and transferred to a 30% sucrose solution 4°C for cryoprotection. After 24–72 hours, brains were flash frozen in isopentane placed in dry ice. For long-term storage, brains were stored in a −80°C freezer. Brains were serially sectioned into four pools at 40 µm-thick coronal sections on a cryostat (Leica 3050S) at −20°C. Sections were placed in 1X PBS solution prior to immunohistochemistry.

### Immunohistochemistry

Sections were washed in PBS and then 0.2% Tween in PBS (PBST), prior to a two-hour blocking incubation in 2% donkey normal serum (DNS) in PBST. Sections were then transferred to the primary antibody (1:1000 dilution in 2% DNS in PBST) for an overnight incubation of 17 hours at room temperature (RT; 21°C). The following primary antibodies were used for staining: MCH (rabbit anti-MCH, H-070-47, Phoenix Pharmaceuticals; RRID:AB_2722682), CART (rabbit anti-CART, H-003-61, Phoenix Pharmaceuticals; RRID:AB_2922768), GFP (chicken anti-GFP, A10262, Invitrogen; RRID:AB_2534023), and NK3R (rabbit anti-NK3R, Novus Biologicals, NB300, RRID:AB_10003270). After incubation, sections were washed in five exchanges of PBST (ten minutes each) and transferred to secondary antibody incubation (1:500 dilution in 2% DNS in PBST) for two hours at RT. The following secondary antibodies were used: donkey anti-rabbit conjugated to Alexa 488 (ab150073, Abcam; RRID:AB_2636877), donkey anti-rabbit conjugated to Alexa 594 (ab150076, Abcam; RRID:AB_2782993), and goat anti-chicken conjugated to Alexa 488 (ab150169, Abcam; RRID:AB_2636803). After incubation, sections were washed first in five washes of PBST and then in five washes of PBS (ten minutes each) prior to mounting onto glass microscope slides with VECTASHIELD HardSet Antifade Medium with DAPI (H-1500-10, Vector Laboratories) or with ProLong Gold Antifade Mountant with DAPI (P36931; Invitrogen). NK3R immunostaining protocol is as described in Fujita et al., 2021. Slides were cured overnight prior to imaging.

### Nissl stain

As previously described (Miller et al., 2024; Negishi et al., 2020), sections adjacent to experimental tissue of virally labelled fibers from DIO-EYFP, CV-Con/Fon-GFP, and Con/Foff injected mice containing the septal complex were mounted on Superfrost Plus microscope slides (Fisher Scientific). The mounted tissue was dehydrated in washes (3 minutes each) of 50%, 70%, 95%, and 100% ethanol, delipidated in xylene, then rehydrated in decreasing concentrations of ethanol, and rinsed in distilled water. Next, the tissue was stained with a 0.25% (w/v) thionine dye solution (pH 4.5) at RT for about five seconds. The reaction was quenched and washed in distilled water, dehydrated in increasing concentrations of ethanol, then cleared in xylene. The sections were immediately coverslipped using Epredia Mounting Medium (Fisher Scientific).

### Fluorescence *in situ* hybridization (FISH) experiments

For FISH experiments, 8-week-old *Tacr3*-Flpo animals were used. For tissue preparation, mice were anesthetized with isoflurane and brains were dissected and flash frozen onto an aluminum sheet in dry ice. Brains were then imbedded into an optimal cutting temperature (OCT) compound block, prior to sectioning. Embedded brains were cryosectioned (Leica 3050S) at 14 µm-thick sections and mounted directly onto Superfrost Plus microscope slides. Collected sections were fixed in 4% paraformaldehyde (PFA) for 15 min at 4°C and dehydrated with increasing concentrations of ethanol (50, 70, 100%). For all experiments, we adhered to protocols provided with the RNAscope Fluorescent Multiplex Reagent Kit v1 by Advanced Cell Diagnostics (ACD). RNAscope probes were designed and validated by ACD and include: *Tacr3* (481671, ACD) and *Flpo* (520791, ACD). Slides were coverslipped with ProLong Gold Antifade Mountant with DAPI.

### Epifluorescence, confocal, and brightfield imaging

Tiled and stitched 10X magnification (0.45 numerical aperture, NA) images of brain sections were obtained using an epifluorescence microscope (BZ-X800 Keyence). Selected areas were then imaged at either (magnification/ NA) 20X/0.75, 40X/1.30, or 100X/1.40 in 1 µm thick stacks using a confocal microscope (Leica SP8). Epifluorescence imaging was performed using consistent brightness and contrast settings; confocal images were taking using consistent laser power settings. For ConVERGD viral anatomical tracing experiments, Injection sites were initially surveyed at 10X magnification using a BZ-X800 Keyence for injection site imaging. After confirmation, 20x/0.80 Plan Apo images (5 µm thick stacks) were taken using the THUNDER Imager Tissue epifluorescent microscope (Leica) and large volume computationally cleared (Felts et al., 2020) to minimize background signal throughout the field of view. Computational clearing was conducted consistently, based on the refractive index of the mounting media used (Vectashield DAPI in all sections). Brightfield images of adjacent Nissl-stained sections were imaged on the BZ-X800 Keyence. Additionally, images within experiment were processed to increase brightness and contrast consistently *post hoc* using Adobe Photoshop CS6.

### Cell quantification analysis

Quantification of FISH colocalization was conducted using a MATLAB script to make a composite of mask of mRNA expressing cells in 40X confocal images based on total mRNA expressing cells. Next, a custom CellProfiler pipeline was used to quantify mRNA fluorescence and colocalization of fluorescence in each cell. Immunostaining quantification was conducted manually using the Cell Counter plug-in for ImageJ (Kurt De Vos, University of Sheffield) using stitched 10X photomicrographs (Keyence) or 40X photomicrographs (confocal). For NK3R quantification, cells were classified as NK3R positive if immunoreactivity was continuously or near-continuously distributed around the soma in confocal photomicrographs at a magnification of 40X. In parallel with all quantification, photomicrographs were registered to Bregma values or distance from Bregma (mm) using Paxinos and Franklin’s the Mouse Brain in Stereotaxic Coordinates (3^rd^ Edition) (Paxinos and Franklin 2012). Off-target or incomplete viral injections were removed from cell body and fiber analysis. Graphs were created using GraphPad Prism (version 10) or Microsoft Excel.

### Fiber analysis

Qualitative assessment of GFP+ and EYFP+ fiber density was performed across brain regions in three viral conditions: total MCH (AAV2-Ef1a-DIO-EYFP in *Pmch*-Cre mice; N=3), Type I MCH (AAV2/8-hSyn-CV-Con/Fon-GFP in *Pmch-*Cre; *Tacr3-*Flpo mice; N=3), and Type II MCH (AAV2/8-hSyn-CV-Con/Foff-GFP in *Pmch-*Cre; *Tacr3-*Flpo mice; N=3). We collected 40 μm-thick coronal sections in four series spanning the antero-posterior extent of the brain (distance from Bregma between +1.98 mm through −6.96 mm). Brain regions were identified with reference to the Allen Adult Mouse Brain Atlas (http://atlas.brain-map.org/). Areas of interest were first established by assessing fiber presence in the DIO-EYFP condition.

Subregions where no EYFP+ fibers were observed in the *Pmch-*Cre+DIO-EYFP condition were omitted from scoring. For each region of interest, multiple sections were examined across the anteroposterior extent of the subregion, and the most representative section was selected for scoring in all three conditions. Fiber density was assessed on a five-point scale: -- (absent), + (sparse), ++ (moderate), +++ (dense), and ++++ (very dense) (representative examples shown in Figure 19). Scores were compared within each condition to confirm consistency. Fiber density was then compared across total MCH, CV-Con/Fon, and CV-Con/Foff, conditions to identify subregions of preferential innervation by each subpopulation.

### Fiber tracing and mapping

For fiber tracing, adjacent Nissl sections were assigned to equivalent coronal Allen Reference Atlas (ARA) level (Dong et al., 2008) and parcellated into subregions according to cell morphology and proximity to white matter and ventricles using Affinity Designer 2. Next, adjacent experimental sections containing fluorescently labeled fibers were overlayed onto the parcellated Nissl image and aligned. Using a graphic drawing tablet (Wacom), fibers were manually traced from experimental images. After tracing, fibers were mapped onto corresponding ARA levels to create standardized fiber maps. This technique was adapted from (Bono et al., 2025; Miller et al., 2024; Negishi et al., 2020; Payant et al., 2024).

### Stimulated Emission Depletion (STED) super-resolution microscopy

Tissue preparation and immunostaining for STED microscopy: For STED microscopy, following cryoprotection, brains of *Pmch*-Cre;*Tacr3*-FlpO mice injected with either AAV2/8-hSyn-CV-Con/Fon-GFP or AAV2/8-hSyn-CV-Con/Foff-GFP were embedded in OCT in cryomolds over freezing isopentane. Coronal brain cryosections (6 μm) were directly mounted onto #1.5 gelatin coated glass coverslips (Electron Microscopy Sciences). Sections were washed three times in PBS (10 min each) and then blocked in 5% DNS in 0.5% Triton-X in PBS for one hour before incubation in the following primary antibodies diluted in blocking solution at a 1:1000 concentration overnight (RT): chicken anti-VGLUT2 (135 409, Synaptic Systems, RRID: AB_2943526) and either rabbit anti-MCH, or rabbit anti-CART. Tissues were subsequently washed with PBS three times (10 min each) then incubated for two hours (RT) with the following secondary antibodies diluted in blocking solution at a 1:200 concentration: goat anti-rabbit IgG STAR ORANGE (STORANGE-1002-500UG, Abberior) and goat anti-chicken STAR RED (STRED-1005-500UG, Abberior).

After three subsequent PBS washes, the tissues were directly mounted onto glass slides and coverslipped with Prolong Diamond mounting media without DAPI (P36970, Invitrogen). STED photomicrographs were taken using an Abberior Instruments Expert Line STED microscope equipped with a 775 nm (for STAR ORANGE and STAR RED) depletion laser and an Olympus 100x/1.40 UPLANSAPO oil immersion objective, through the University of Connecticut Advanced Light Microscopy Facility (RRID:SCR_027547).

## RESULTS

### Generation and validation of intersectional genetic tools for access to MCH subpopulations

A considerable barrier in investigating MCH subpopulations has been the lack of specific genetic models available to selectively manipulate the spatially intermixed, yet genetically distinct MCH subpopulations. To overcome this, we developed parallel intersectional genetic strategies utilizing a newly developed knock-in *Tacr3*-Flpo mouse line, in which Flpo recombinase expression is driven by the endogenous *Tacr3* promoter **(Fig. 1A).** We use this mouse line to leverage *Tacr3* expression as a molecular toggle to intersectionally divide the MCH population. For this study, we operationally defined Type I MCH neurons as *Pmch+/Tacr3+* and Type II MCH neurons as *Pmch+/Tacr3-.* As the reliability of our system depends on this previously uncharacterized *Tacr3*-Flpo mouse line, we first investigated the specificity and penetrance of the Flpo expression relative to endogenous *Tacr3*.

**Figure 1:**
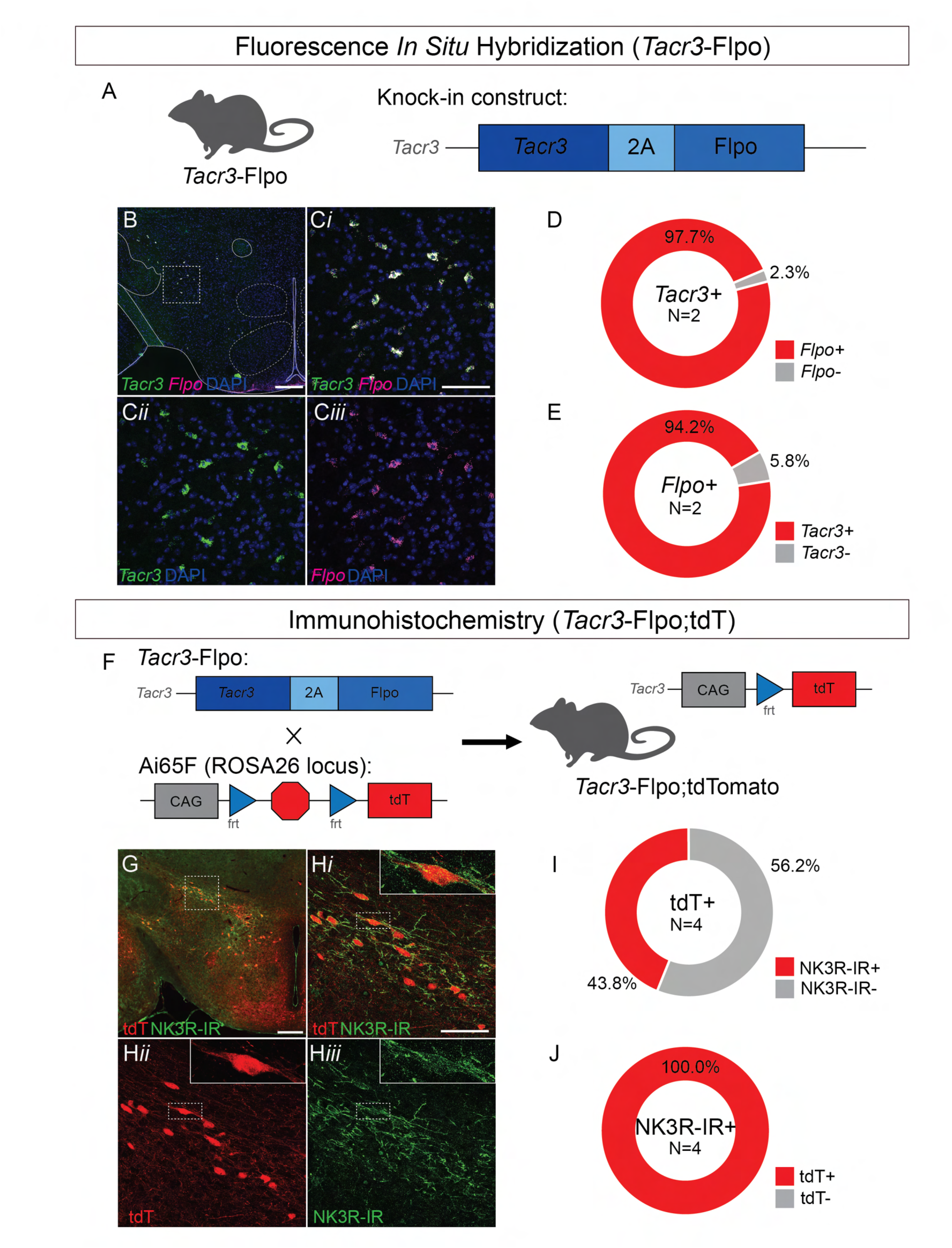
Fluorescence *in situ* hybridization and immunohistochemical validation of the *Tacr3*-Flpo line. **A)** Schematic depicting the generation of the knock-in *Tacr3*-Flpo mouse line. **B-C)** Representative 10X epifluorescent photomicrograph **(B)** and 40X confocal photomicrograph **(C)** taken from outlined area of the LHA of *Tacr3*-Flpo animals showing colocalization (***i***) of *Tacr3* mRNA (green; ***ii***) and *Flpo* mRNA hybridization (magenta; ***iii***). **D)** Donut plot indicating *Flpo* mRNA specificity in total *Tacr3* mRNA expressing cells in the LHA of *Tacr3-*Flpo mice (275 cells, N=2). **E)** Donut plot indicating *Tacr3* mRNA penetrance in total *Flpo* expressing cells in the LHA of *Tacr3*-Flpo mice (n=265 cells; N=2). **F)** Schematic depicting cross of heterozygous *Tacr3-*Flpo animals with homozygous Ai65F (Daigle et al., 2018) to generate a *Tacr3*-Flpo;tdTomato (tdT) animal. **G)** Representative 10X epifluorescence from the LHA of *Tacr3*-Flpo;tdT animals showing colocalization of NK3R immunoreactivity (NK3R-IR; green) and tdTomato (tdT; red). **H**) Confocal photomicrographs from outlined area in (G) showing colocalization (***i***) of tdTomato (red; ***ii***) and NK3R-IR (green; ***iii***). 100X Higher resolution confocal photomicrograph within insets (***i***–***iii***) show colocalization of tdT and NK3R at individual cells. (**I**) Donut plot indicating NK3R-expressing tdT+ cells in the LHA (n=737 tdT+ cells; N=4). (**J**) Donut plot indicating tdT-expressing NK3R-IR+ cells in the LHA (n=326 NK3R-IR+ cells; N=4). 10X scale bars: 300 µM. 40X scale bars: 100µM.

Using fluorescence *in situ* hybridization (FISH), we first probed for *Tacr3* and *Flpo* mRNA in the LHA (**Fig. 1B**, **1C**) and then used unbiased cell counting pipelines and identified that *Flpo* mRNA was co-expressed in 97.7% (n=275 cells, N=2 animals) of *Tacr3* mRNA expressing cells in the LHA (**Fig. 1D**), confirming high specificity of the knock-in allele specific to *Tacr3* cells. We next assessed penetrance by quantifying the proportion of *Tacr3*+ cells that co-expressed *Flpo* and identified that *Flpo* mRNA is present in 94.2% of *Tacr3*+ cells (n=265 cells; N=2) **(Fig. 1E)**. We also observed high specificity and penetrance in neighboring subregions such as the arcuate nucleus, ventral tegmental area, and interpeduncular nucleus (unpublished data), consistent with previous work in the rat brain showing immunostaining for neurokinin-3 receptor (NK3R), the protein product of *Tacr3* (Ding et al., 1996).

Next, to visualize Flpo expression in the *Tacr3*-Flpo line, we crossed these mice into a Flpo-dependent tdTomato (tdT; Ai65F) reporter line (Daigle et al., 2018), thus generating a *Tacr3*-Flpo;tdT mouse line. **(Fig. 1F)**. We next assessed if NK3R is expressed in tdT-labeled cell bodies in these mice. NK3R immunostaining in hypothalamic sections of the *Tacr3*-Flpo;tdT mouse revealed diffuse immunoreactivity (IR) localized to the cell body surface and somatic processes, consistent with previous reports in the LHA **(Fig. 1G, 1H)** (Fujita et al., 2021; Miller et al., 2024). Only 43.8 ± 3.0% (mean ± SEM; 737 tdT+ cells; N=4) of tdT+ cell bodies exhibited NK3R-IR (**Fig. 1I**), as expected, given that mRNA expression does not always correspond to detectable protein levels. Additionally, the Ai65F reporter may serve as a fate map, in expressing tdT fluorescence in cells that may transiently express *Tacr3* during development, even when *Tacr3* expression is no longer present in the adult. Importantly, all NK3R-IR cell bodies colocalized with tdT fluorescence (100% ± 0.0%; 326 NK3R-IR+ cells; N=4) **(Fig. 1J)**, confirming that the *Tacr3*-Flpo line reports the full spectrum of NK3R-expressing neurons. Together, these FISH and immunohistochemical validation experiments established the *Tacr3*-Flpo line as a reliable tool in gaining genetic access to and manipulation of *Tacr3*-expressing MCH neurons.

### Immunohistochemical characterization of MCH subpopulations in the *Tacr3-*Flpo;tdT mouse line

We next asked if Type I and Type II MCH subpopulations could be identified using the *Tacr3*-Flpo;tdT mouse line. Although spatially intermixed, previous work in rats and mice indicates that Type I and Type II MCH neurons exhibit distinct spatial distribution profiles across the LHA, where Type I neurons are enriched in anterior and medial regions, while Type II neurons are denser more posteriorly and laterally (Croizier et al., 2010; Cvetkovic et al., 2004; Miller et al., 2024; Wang et al., 2021). To determine whether this distribution was recapitulated in our *Tacr3*-Flpo;tdT reporter line, we stained for the MCH protein throughout the male and female LHA **(Fig. 2).** We classified Type I MCH neurons as MCH-IR+/tdT+ and Type II MCH neurons as MCH-IR+/tdT-. Consistent with previous reports, MCH-IR and tdT+ colocalization revealed distinct patterns along the anteroposterior extent of the hypothalamus **(Fig. 2A-D).** A larger proportion of MCH-IR+/tdT+ neurons were observed in anterior sections of the LHA (70.9 ± 2.7%; -1.34 mm from bregma; N=5) compared to more posterior sections (47.9 ± 4.0%; -1.70 mm from bregma; N=5) **(Fig. 2E)**. We also observed some tdT+ neurons in the LHA that lacked MCH-IR; indicating that *Tacr3* expression is not restricted to MCH-IR cell bodies. Qualitatively, these few MCH-IR−/tdT+ cells appeared smaller in size compared to MCH-IR+/tdT+ cells **(Fig. 2B)**. Overall, 60.3 ± 2.7% of total MCH-IR neurons expressed tdT+ (4874 total MCH-IR+ cells; N=5) **(Fig. 2F)**, consistent with our single-cell RNA sequencing data.

**Figure 2:**
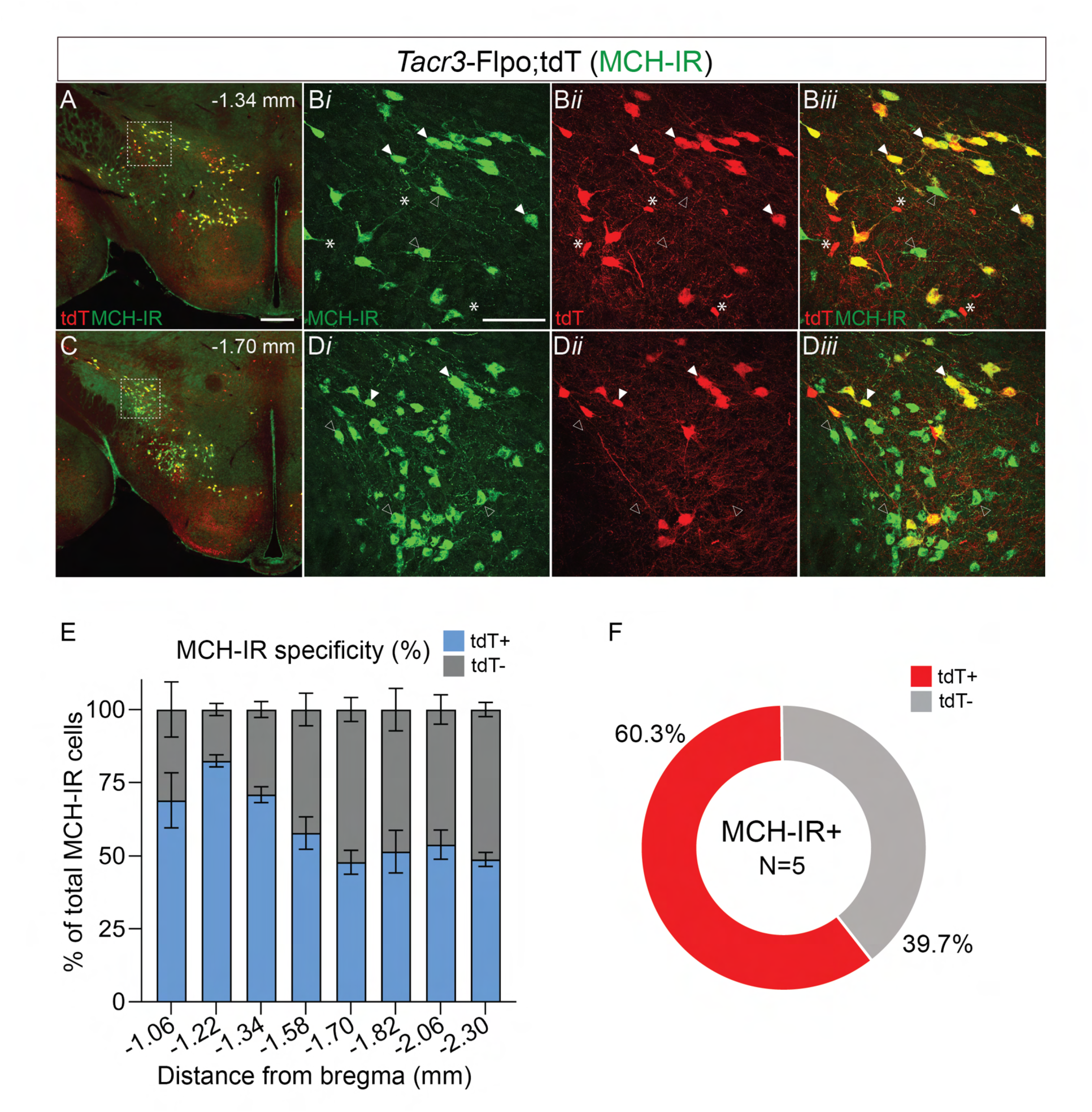
Characterization of *Tacr3-*Flpo driven tdTomato expression in MCH-immunoreactive cells in the hypothalamus. A-D) Representative 10X epifluorescent merged photomicrograph of tdT expression (red) and MCH-IR (green) at -1.34 mm **(A)** and -1.70 mm **(D)** of *Tacr3*-Flpo; tdT mice. Confocal photomicrographs (40X) **(B, D)** from outlined area in **(A)** and **(D)** showing MCH-IR (green; ***i***), tdT (red; ***ii***), and colocalization (***iii***). White arrows point towards representative Type I cells (MCH-IR+/tdT+). Outlined arrows point towards representative Type II cells (MCH-IR+/tdT-). Asterisks indicate representative cells that are MCH-IR-/tdT+. **E)** % of tdT+ (blue) and tdT- (gray) MCH-IR+ cell bodies throughout the LHA (N=5 mice). Error bars represent standard error of means (SEM). **F)** Donut plot indicating tdT+ specificity in MCH-IR+ cell bodies in *Tacr3*-Flpo; tdT mice (4874 total MCH-IR+ cells; N=5). MCH-IR/tdT+ specificity across the anteroposterior axis (distance from bregma): 68.9 ± 9.4% (−1.06 mm), 82.4 ± 2.1% (−1.22 mm), 70.9 ± 2.7% (−1.34 mm), 57.8 ± 5.5% (−1.58 mm), 47.9 ± 4.0% (−1.70 mm), 51.5 ± 7.2% (−1.82 mm), 53.9 ± 5.0% (−2.06 mm), and 48.8 ± 2.4% (−2.30 mm) (N=5). 10X scale bars: 300 µM. 40X scale bars: 100µM.

We next examined protein expression of the Type I MCH neuron marker, cocaine and amphetamine regulated transcript (CART), in LHA tdT+ neurons **(Fig. 3A-D).** Consistent with the distribution of Type I MCH neurons, CART-IR expression was most abundant in anterior sections of the LHA (94.1 ± 0.7%; -1.34 mm from bregma; N=5) compared to posterior sections (89.7 ± 2.6%; -1.70 mm from bregma; N=5) (**Fig. 3E).** Overall, 89.7 ± 1.9% of total CART-IR cells expressed tdT (3072 total CART-IR cells; N=5), confirming high fidelity between the *Tacr3*-Flpo reporter and CART expression **(Fig. 3F).** However, we also observed a small proportion of CART-IR neurons that did not express tdT (10.1 ± 1.9%), consistent with single-cell RNA sequencing of the hypothalamus, which indicates the presence of *Tacr3*-independent CART populations that exist in the LHA (Mickelsen et al., 2019). Together, these data validate the *Tacr3*-Flpo line as a reliable tool for distinguishing Type I and Type II MCH neurons, thus enabling subpopulation-specific investigation of MCH circuitry.

**Figure 3:**
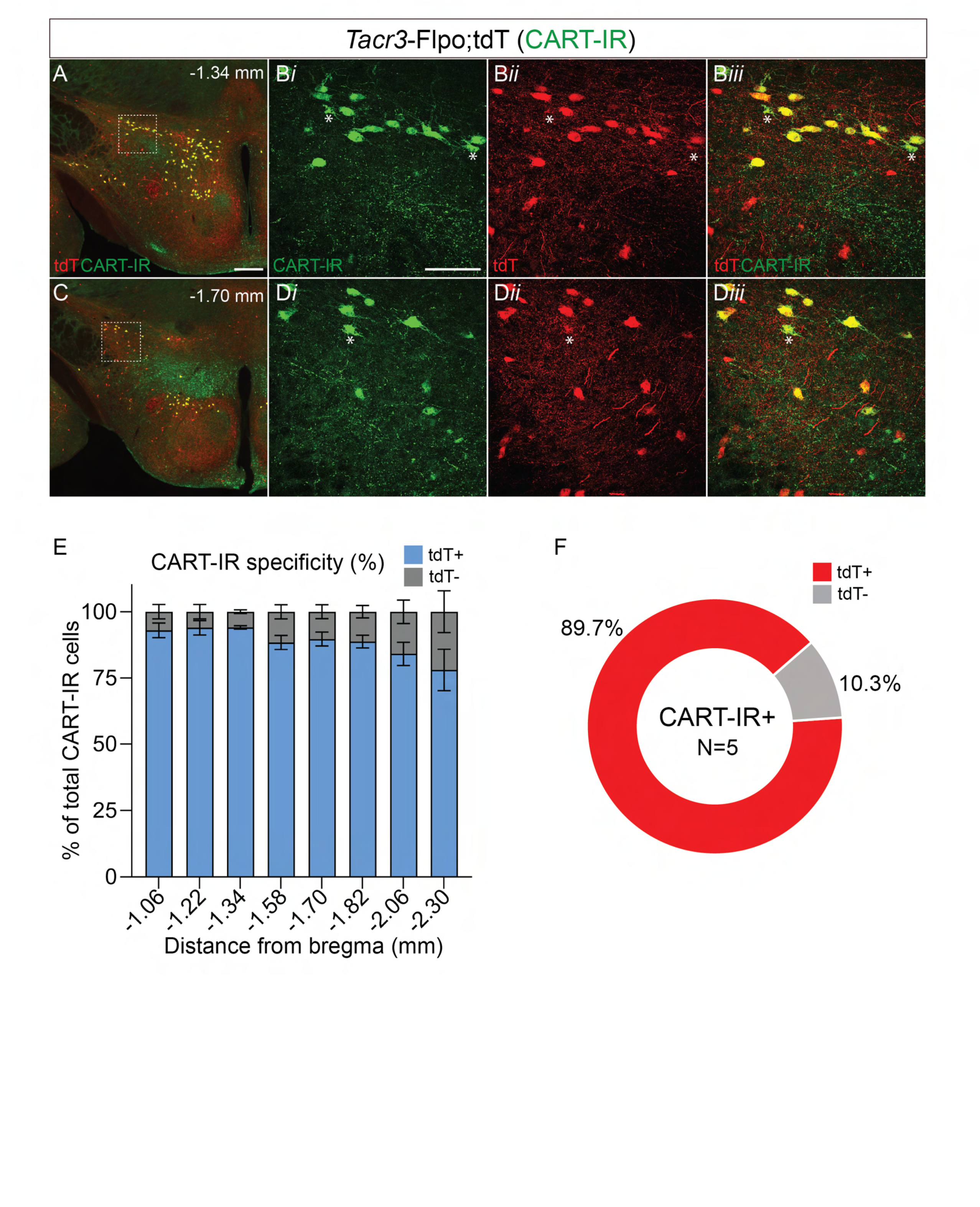
Characterization of *Tacr3-*Flpo driven tdTomato expression in CART-immunoreactive cells in the hypothalamus. A-D) Representative 10X epifluorescent merged photomicrograph of tdT expression (red) and CART-IR (green) at -1.34 mm **(A)** and -1.70 mm **(C)** of *Tacr3*-Flpo; tdT mice. Confocal photomicrographs **(B, D)** from outlined area in **(A)** and **(C)** showing CART-IR (green; ***i***), tdT (red; ***ii***), and colocalization **(*iii*)**. Asterisks indicate representative cells that are CART-IR+/tdT-. **E)** Quantification of tdT+ (blue) and tdT- (gray) CART-IR cell bodies throughout the LHA (N=5 mice). Error bars represent standard error of means (SEM). **F)** Donut plot indicating tdT+ specificity in CART-IR+ cell bodies in *Tacr3*-Flpo; tdT mice (3072 total CART-IR+ cells; N=5). CART-IR/tdT+ specificity across the anteroposterior axis (distance from bregma): 93.0 ± 2.7% (−1.06 mm), 94.0 ± 2.7% (−1.22 mm), 94.1 ± 0.7% (−1.34 mm), 88.4 ± 2.7% (−1.58 mm), 89.7 ± 2.6% (−1.70 mm), 88.8 ± 2.4% (−1.82 mm), 84.1 ± 4.4% (−2.06 mm), and 78.1 ± 7.8% (−2.30 mm) (N=5).10X scale bars: 300 µM. 40X scale bars: 100 µM.

### Identification of Type I MCH projections using an intersectional reporter mouse line

To gain genetic access to MCH subpopulations, we crossed heterozygous *Pmch*-Cre (Kong et al., 2010) and *Tacr3*-Flpo mice to generate a dual-recombinase *Pmch*-Cre;*Tacr3*-Flpo mouse line which served as a foundation in our intersectional reporter and viral strategies **(Fig. 4A)**. First, for our intersectional reporter strategy, we crossed this *Pmch*-Cre;*Tacr3*-Flpo mouse line to a RC::FLTG dual recombinase reporter line (Plummer et al., 2015) **(Fig. 4B)** to generate a triple mutant *Pmch*-Cre; *Tacr3*-Flpo; RC::FLTG mouse. In this line, successful dual recombination of Cre AND Flpo drives GFP fluorescence in intersectionally-targeted cells and their axonal projections (Plummer et al., 2015), which we hypothesized to correspond to Type I MCH neurons.

**Figure 4:**
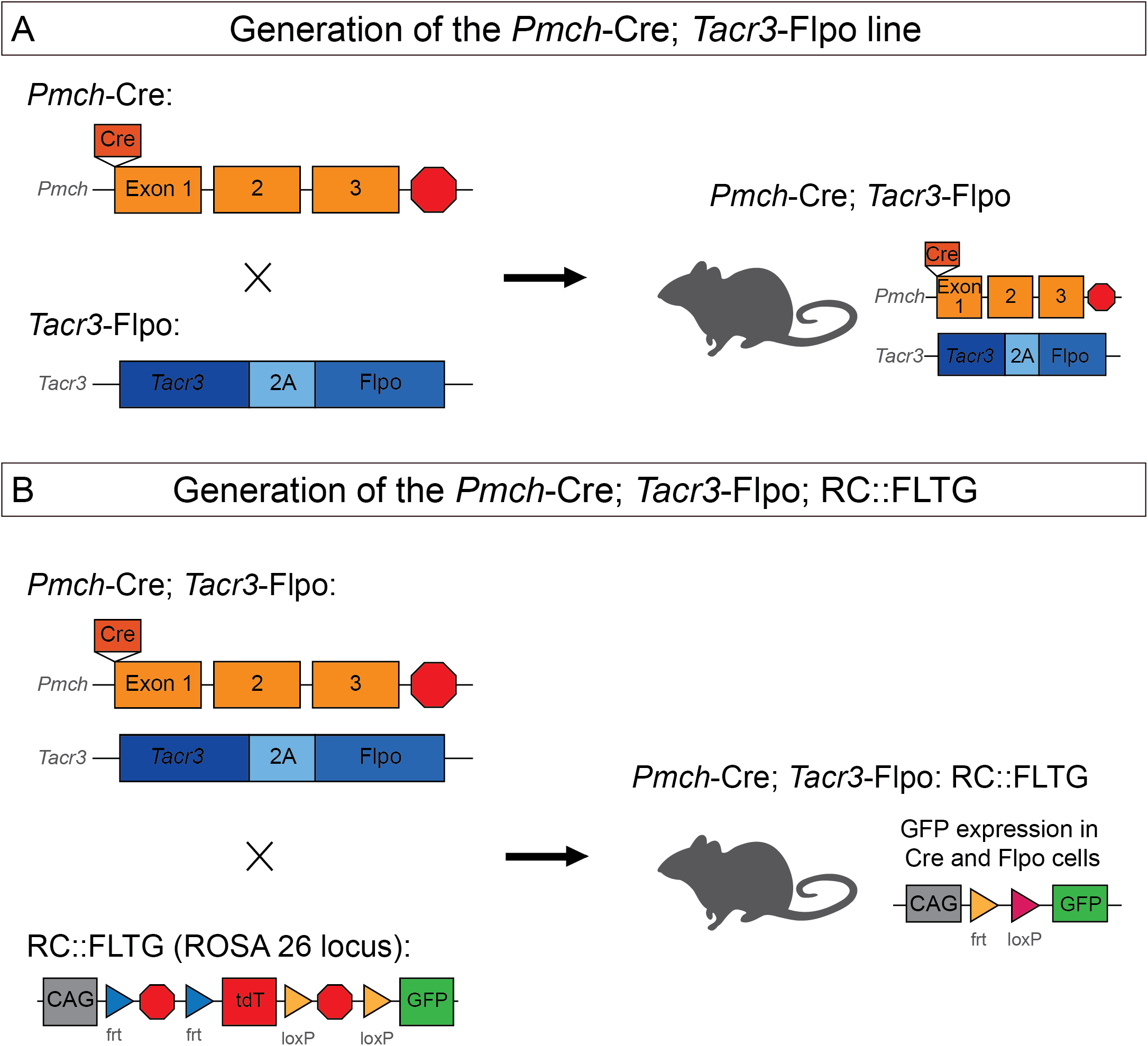
Generation of intersectional mouse models for genetic access to MCH subpopulations. **A)** Schematic depicting breeding of heterozygous BAC transgenic *Pmch*-Cre (Kong et al., 2010) and heterozygous knock-in *Tacr3*-Flpo mice to generate dual recombinase *Pmch*-Cre; *Tacr3*-Flpo line. The *Pmch*-Cre mouse line has been previously validated for Cre recombinase specificity (Beekly et al., 2020; Mickelsen et al., 2017; Miller et al., 2024). Schematic of BAC transgene adapted from Kong et al., 2010. **B)** For intersectional reporting of Type I MCH neurons, we crossed our dual recombinase *Pmch*-Cre;*Tacr3*-Flpo mouse **(A)** to a dual recombinase reporter RC::FLTG (Plummer et al., 2015) to generate a triple mutant *Pmch*-Cre; *Tacr3*-Flpo; RC::FLTG mouse for the selective expression of GFP in *Cre* AND *Flpo* expressing neurons. Schematic of RC::FLTG ROSA26 locus adapted from Plummer et al., 2015.

Before examining Type I MCH subcircuits, we first assessed if this triple mutant line specifically reports Type I MCH cell bodies in the LHA. GFP immunostaining was used to amplify endogenous GFP fluorescence. GFP-IR showed high specificity to MCH-IR neurons throughout the LHA **(Fig. 5A-B**). As expected, a higher proportion of MCH-IR+/GFP-IR+ neurons were observed in anterior sections of the LHA (56.0 ± 6.5%; -1.34 mm from bregma; N=5) compared to posterior sections (30.6 ± 6.0%; -1.70 mm from bregma; N=5) **(Fig. 5C).** Overall, 98.0 ± 0.6% of total GFP-IR+ neurons expressed MCH-IR (1875 total GFP-IR cells; N=5) **(Fig. 5D)** and 42.3 ± 3.6% of total MCH-IR neurons expressed GFP-IR (4479 total MCH-IR cells; N=5) **(Fig. 5E).**

**Figure 5:**
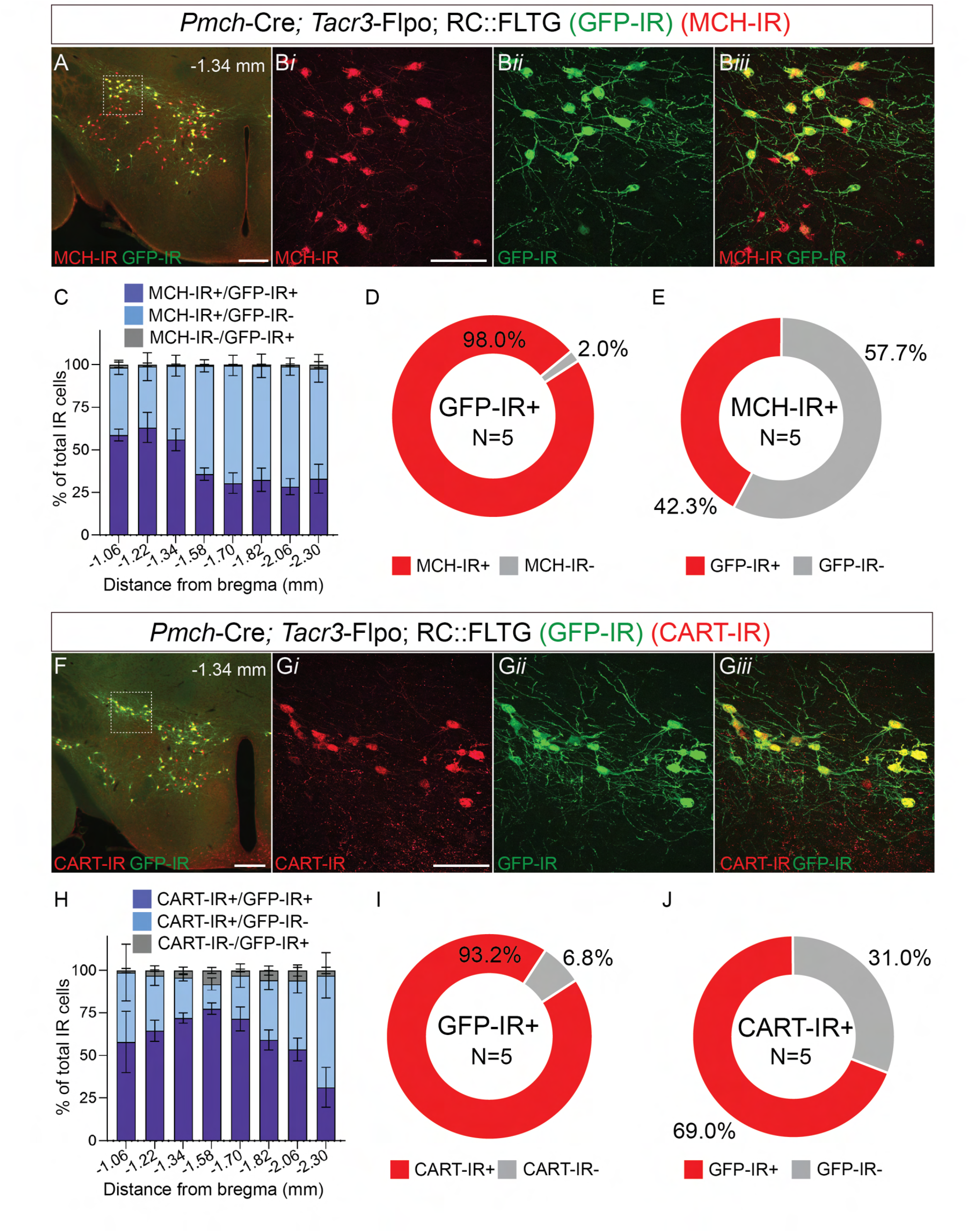
Intersectional reporter mouse line selectively labels Type I MCH neurons in the LHA. **A)** Representative merged 10X epifluorescent photomicrograph of GFP-IR (green) and MCH-IR (red) at -1.34 mm from bregma in the LHA of a *Pmch*-Cre; *Tacr3*-Flpo; RC::FLTG mouse. GFP-IR was used to boost endogenous GFP signal from the RC::FLTG reporter. **B)** Confocal photomicrographs (40X) from outlined area in (A) showing MCH-IR (red; ***i***), GFP-IR (green; ***ii***), and colocalization (***iii***). **C)** Representation of proportion of MCH-IR+/GFP-IR+ (dark blue), MCH-IR+/GFP-IR- (light blue) and MCH-IR-/GFP-IR+ (gray) cells throughout the LHA measured by distance from bregma of *Pmch*-Cre;*Tacr3*-Flpo; RC::FLTG (N=5). Error bars represent ± SEM. MCH-IR+/GFP-IR+ across the anteroposterior axis (distance from bregma): 58.8 ± 3.5% (−1.06 mm), 63.2 ± 8.9% (−1.22 mm), 56.0 ± 6.5% (−1.34 mm), 35.9 ± 3.6% (−1.58 mm), 30.6 ± 6.0% (−1.70 mm), 32.5 ± 6.8% (−1.82 mm), 28.5 ± 4.8% (−2.06 mm), and 33.1 ± 8.5% (−2.30 mm). MCH-IR+/GFP-IR-across the anteroposterior axis (distance from bregma): 39.7 ± 4.1% (−1.06 mm), 35.7 ± 8.2% (−1.22 mm), 43.5 ± 6.1% (−1.34 mm), 63.5 ± 3.6% (−1.58 mm), 69.0 ± 6.0% (−1.70 mm), 66.9 ± 6.8% (−1.82 mm), 70.5 ± 5.0% (−2.06 mm), and 64.8 ± 8.1% (−2.30 mm). **D-E)** Donut plots representing GFP-IR penetrance (n=1875 GFP-IR+ cells; N=5) **(D)** and MCH-IR specificity (n= 4479 MCH-IR+ cells; N=5) **(E)** in the LHA of *Pmch*-Cre;*Tacr3*-Flpo; RC::FLTG. **F)** Representative merged 10X epifluorescent photomicrograph of GFP-IR (green) and CART-IR (red) at -1.34 mm from bregma in the LHA of a *Pmch*-Cre; *Tacr3*-Flpo; RC::FLTG mouse. GFP-IR was used to boost endogenous GFP signal from the RC::FLTG reporter. **G)** Confocal photomicrographs (40X) from outlined area in (F) showing CART-IR (red; ***i***), GFP-IR (green; ***ii***), and colocalization (***iii***). **H)** Representation of proportion of CART-IR+/GFP-IR+ (dark blue), CART-IR+/GFP-IR- (light blue) and CART-IR-/GFP-IR+ (gray) cells throughout the LHA measured by distance from bregma of *Pmch*-Cre;*Tacr3*-Flpo; RC::FLTG (N=5). Error bars represent ± SEM. CART-IR+/GFP-IR+ across the anteroposterior axis (distance from bregma): 57.9 ± 17.9% (−1.06 mm), 64.4 ± 6.2% (−1.22 mm), 72.0 ± 3.0% (−1.34 mm), 77.4 ± 3.3% (−1.58 mm), 71.5 ± 7.0% (−1.70 mm), 59.1 ± 5.9% (−1.82 mm), 53.5 ± 6.6% (−2.06 mm), and 31.3 ± 11.7% (−2.30 mm). CART-IR+/GFP-IR-across the anteroposterior axis (distance from bregma): 40.8 ± 16.8% (−1.06 mm), 32.6 ± 5.8% (−1.22 mm), 23.8 ± 1.9% (−1.34 mm), 14.7 ± 3.6% (−1.58 mm), 25.5 ± 6.9% (−1.70 mm), 35.3 ± 5.6% (−1.82 mm), 40.7 ± 7.6% (−2.06 mm), and 65.8 ± 13.3% (−2.30 mm). **I-J)** Donut plots representing GFP-IR penetrance (n=1992 GFP-IR+ cells; N=5) **(I)** and CART-IR specificity (n= 2793 CART-IR+ cells; N=5) **(J)** in the LHA of *Pmch*-Cre;*Tacr3*-Flpo; RC::FLTG. 10X scale bars: 300 µM. 40X scale bars: 100µM.

We next investigated GFP-IR specificity to CART-IR neurons throughout the anteroposterior extent of the LHA **(Fig. 5F-H).** The majority (93.2 ± 2.0%;) of GFP-IR+ cells co-expressed CART-IR (1992 total GFP-IR cells; N=5) thus confirming high specificity to Type I MCH neurons **(Fig. 5I)**. Notably, the RC::FLTG reporter exhibited incomplete penetrance, as only 69.0 ± 3.7%; of CART-IR cells expressed GFP-IR (2793 total CART-IR cells; N=5) **(Fig. 5J).** This was notably lower than the 89.7 ± 1.9% CART-IR+/tdT+ co-localized cells in the *Tacr3*-Flpo; tdT **(Fig. 4)** and suggested that dual recombination at the RC::FLTG locus is likely less efficient than the tdT+ reporter line. Overall, as the *Pmch*-Cre; *Tacr3*-Flpo; RC::FLTG line demonstrated high specificity for Type I MCH neurons, we proceeded to investigate anterograde projection patterns in this mouse.

We next examined GFP-IR axonal fibers in the *Pmch*-Cre; *Tacr3*-Flpo; RC::FLTG mouse. Overall fiber expression was qualitatively dim, even after GFP immunostaining to amplify native GFP fluorescence. We visualized moderate GFP-IR+ fiber labeling in the anterior cingulate area (ACA), lateral septum (LS), and medial septal complex (MSC) **(Fig. 6A-C).** We identified dense GFP-IR+ fiber labeling in the mammillary body (MBO), ventral tegmental area (VTA), and superior colliculus (SC) (**Fig. 6E**–**G**). We found sparse GFP-IR+ fibers in the dorsal hippocampal formation and inferior colliculus **(Fig. 6D, H)**. Notably, fiber expression in the LS and dHPF contrasted with previous reports of dense innervation by total MCH neurons (Bittencourt et al., 1992; Chee et al., 2015; Izawa et al., 2019; Payant et al., 2024), suggesting that these regions may be innervated by *Pmch+/Tacr3-*neurons or Type II MCH neurons.

**Figure 6:**
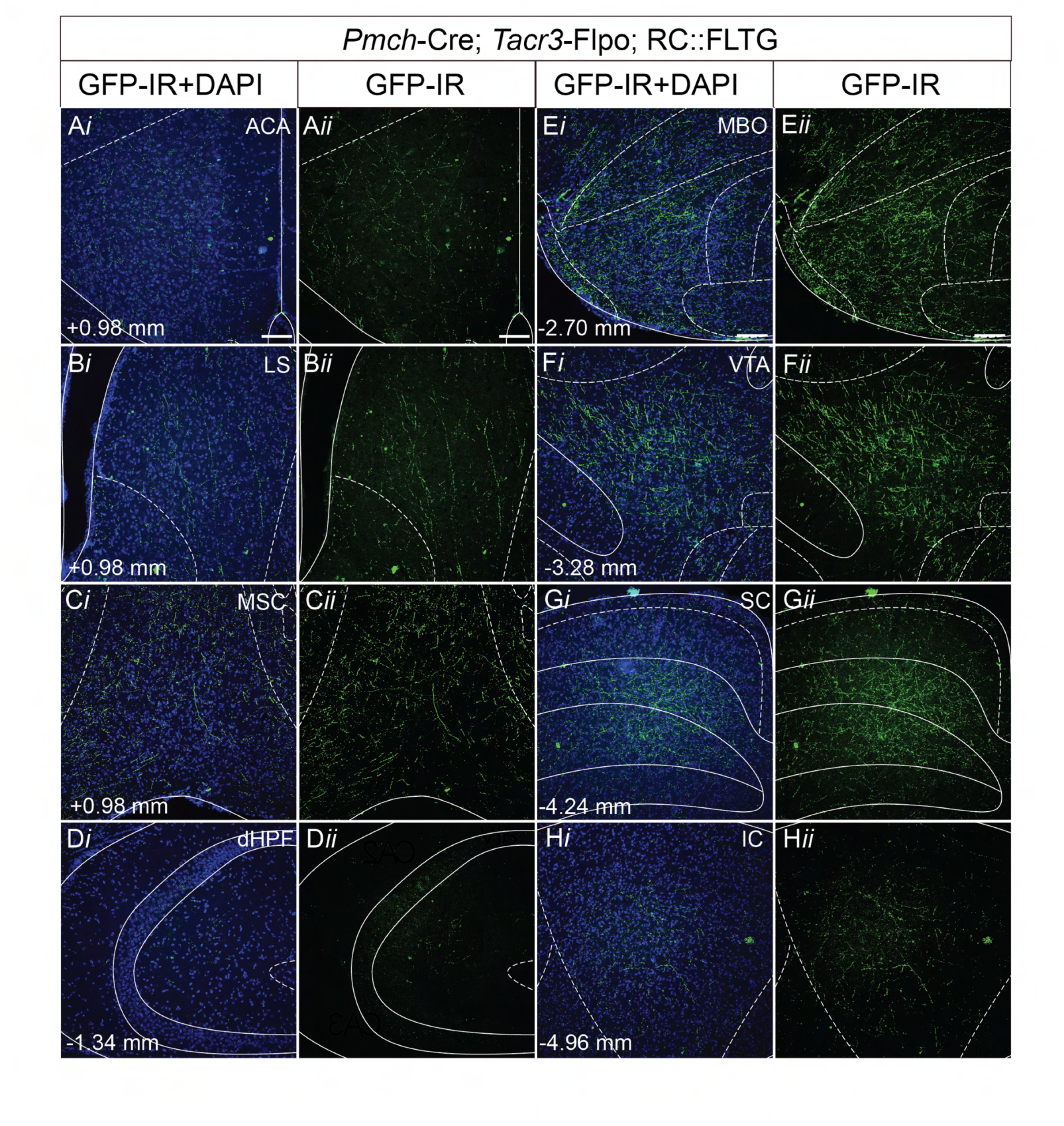
Intersectional reporter labeling reveals Type I MCH anterograde projection patterns. A-H) Representative confocal photomicrographs (20X) of GFP-IR fibers (green) and DAPI nuclear staining (blue, ***i***) and GFP-IR alone **(*ii*)** in the ACA **(A)**, LS **(B)**, MSC **(C)**, dHPF **(D)**, MBO **(E)**, VTA **(F)**, SC **(G)**, and IC **(H)** in *Pmch*-Cre; *Tacr3*-Flpo; RC::FLTG mice (N=5) with corresponding bregma values (lower left). 20X scale bars: 100µM. **Abbreviations:** ACA: anterior cingulate area; dHPF: dorsal hippocampal formation; IC: inferior colliculus; MBO: mammillary body; LS: lateral septum; MSC: medial septum complex; SC: superior colliculus; VTA: ventral tegmental area.

Overall, the *Pmch*-Cre; *Tacr3*-Flpo; RC::FLTG mouse provided highly specific reporting of Type I MCH neurons and foundational understanding of Type I MCH subcircuits. However, this dual recombinase reporter line is limited to the visualization of only Type I MCH cell bodies and axonal projections, thus the use of other intersectional approaches are needed for the identification of Type II MCH neurons and functional comparisons of MCH subpopulations.

### Validation of ConVERGD viruses for intersectional targeting of MCH subpopulations

To more directly compare Type I and Type II MCH neurons, we applied a viral approach using the ConVERGD (Conditional Viral Expression by Ribozyme Guided Degradation) viruses (Hughes et al., 2024) in *Pmch*-Cre;*Tacr3*-Flpo mice (Fig. 4). ConVERGD (CV) adeno-associated viruses (AAV) use a self-cleaving ribozyme controlled by recombinase activity to target neuron subpopulations based on the Boolean logic of Cre and Flpo expression (Hughes et al., 2024).

Using these CV viruses, we devised a comparative approach to assess MCH subpopulation projection patterns **(Fig. 7)**. First, to provide a baseline for comparison, we injected an AAV-Ef1a-DIO-EYFP (or DIO-EYFP) virus into *Pmch*-Cre mice to drive anterograde EYFP expression into all MCH neurons **(Fig. 7A)**. In parallel, we injected *Pmch*-Cre; *Tacr3*-Flpo mice with an AAV-hSyn-CV(Con/Fon)-GFP (or CV-Con/Fon-GFP) virus that drives anterograde GFP expression in cells expressing both Cre and Flpo recombinases to label *Pmch*+/*Tacr3*+ (Type I MCH) neurons **(Fig. 7B).** In another cohort of *Pmch*-Cre;*Tacr3*-Flpo animals, we injected an AAV-hSyn-CV(Con/Foff)-GFP (or CV-Con/Foff-GFP) virus that drives anterograde GFP fluorescence in cells expressing Cre but not Flpo recombinase to label *Pmch*+/*Tacr3*-(Type II MCH) neurons **(Fig. 7C).**

**Figure 7:**
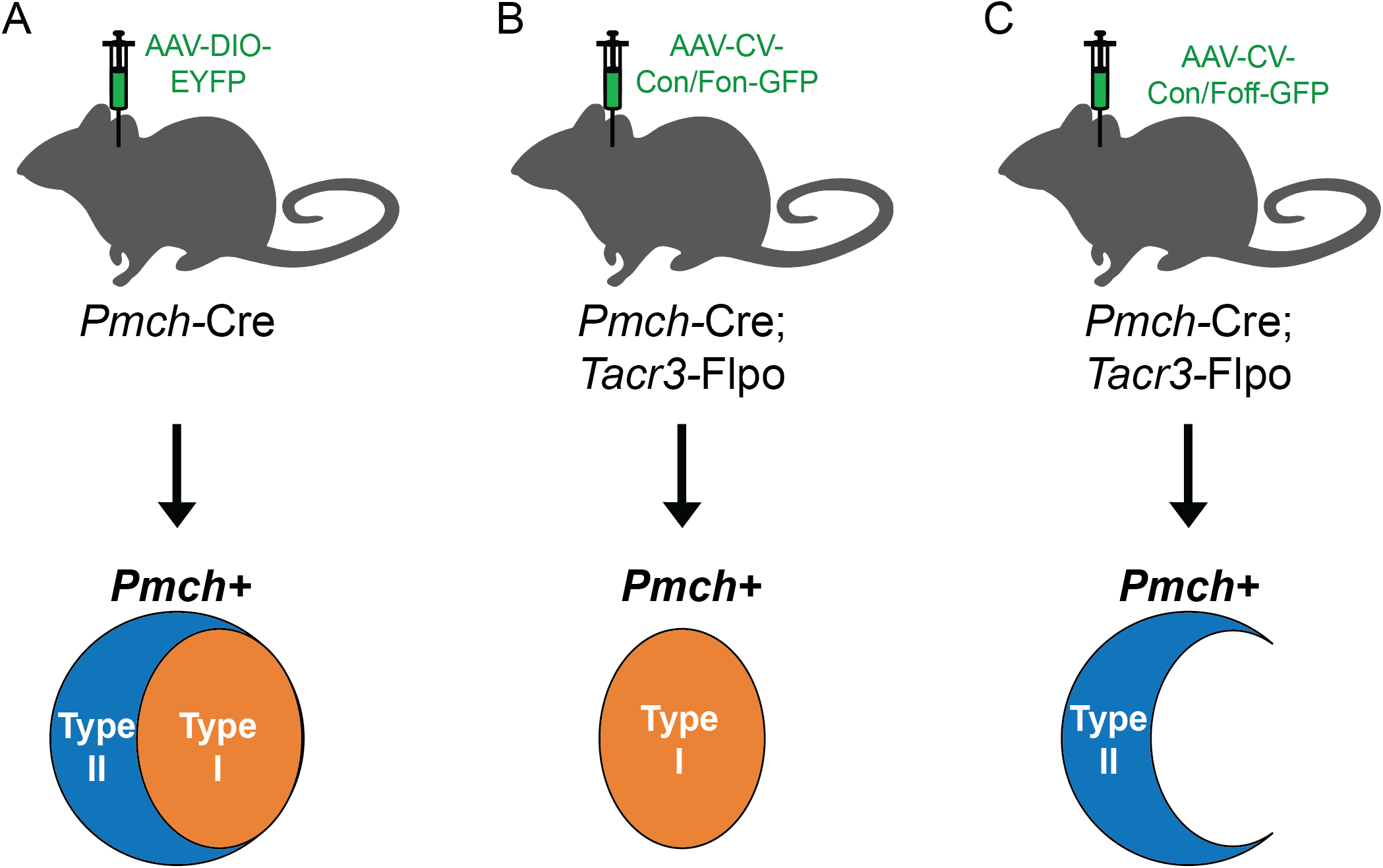
Schematic of intersectional viral targeting strategy. Schematic represents three viral experimental conditions. **A)** *Pmch*-Cre injected with a bilateral injection of AAV2-EF1a-DIO-EYFP will result in anterograde EYFP expression in all *Pmch* cell bodies and axons. **B)** *Pmch*-Cre;*Tacr3*-Flpo injected with a bilateral injection of AAV2/8-hSyn-CV-Con/Fon-GFP will result in anterograde GFP expression in only *Pmch*+/*Tacr3*+ cell bodies and axons. **C)** *Pmch*-Cre;*Tacr3*-Flpo injected with a bilateral injection of AAV2/8-hSyn-CV-Con/Foff-GFP will result in anterograde GFP expression in only *Pmch*+/*Tacr3*-cell bodies and axons.

Prior to injection of ConVERGD viruses in dual recombinase mice, we first assessed viral specificity and potential off-target expression in control animals lacking one or both recombinases. To understand the extent of non-specific ConVERGD viral labeling, we first injected CV-Con/Fon-GFP (**Fig. 8A**) into the LHA of wild type C57BL/6 (**Fig. 8B**), *Pmch*-Cre (**Fig. 8C**), and *Tacr3*-Flpo mice (**Fig. 8D**). We counterstained for MCH and GFP, the latter to enhance virally mediated GFP expression and examined GFP specificity to MCH-IR neurons. In all three control conditions, we observed little to no GFP expression. We next repeated this experiment with the CV-Con/Foff-GFP virus (**Fig. 8E**) and identified little to no GFP expression in the wild type C57BL/6 (**Fig. 8F**) However, we did observe GFP expression in MCH-IR+ neurons in *Pmch*-Cre control mice (**Fig. 8G**). This aligned with our expectations given that the CV-Con/Foff ribozyme only degrades GFP mRNA in the presence of Flpo, and as *Pmch*-Cre mice lack Flpo recombinase, Cre-mediated expression of GFP would proceed without ribozyme-mediated silencing. In *Tacr3*-Flpo controls, we identified limited non-specific viral labeling, consistent with previous reporting (Hughes et al., 2024) **(Fig. 8H).** Together, these results confirmed that CV-Con/Fon-GFP and CV-Con/Foff-GFP viral targeting elicited minimal off-target expression prior to injection in dual recombinase mice.

**Figure 8:**
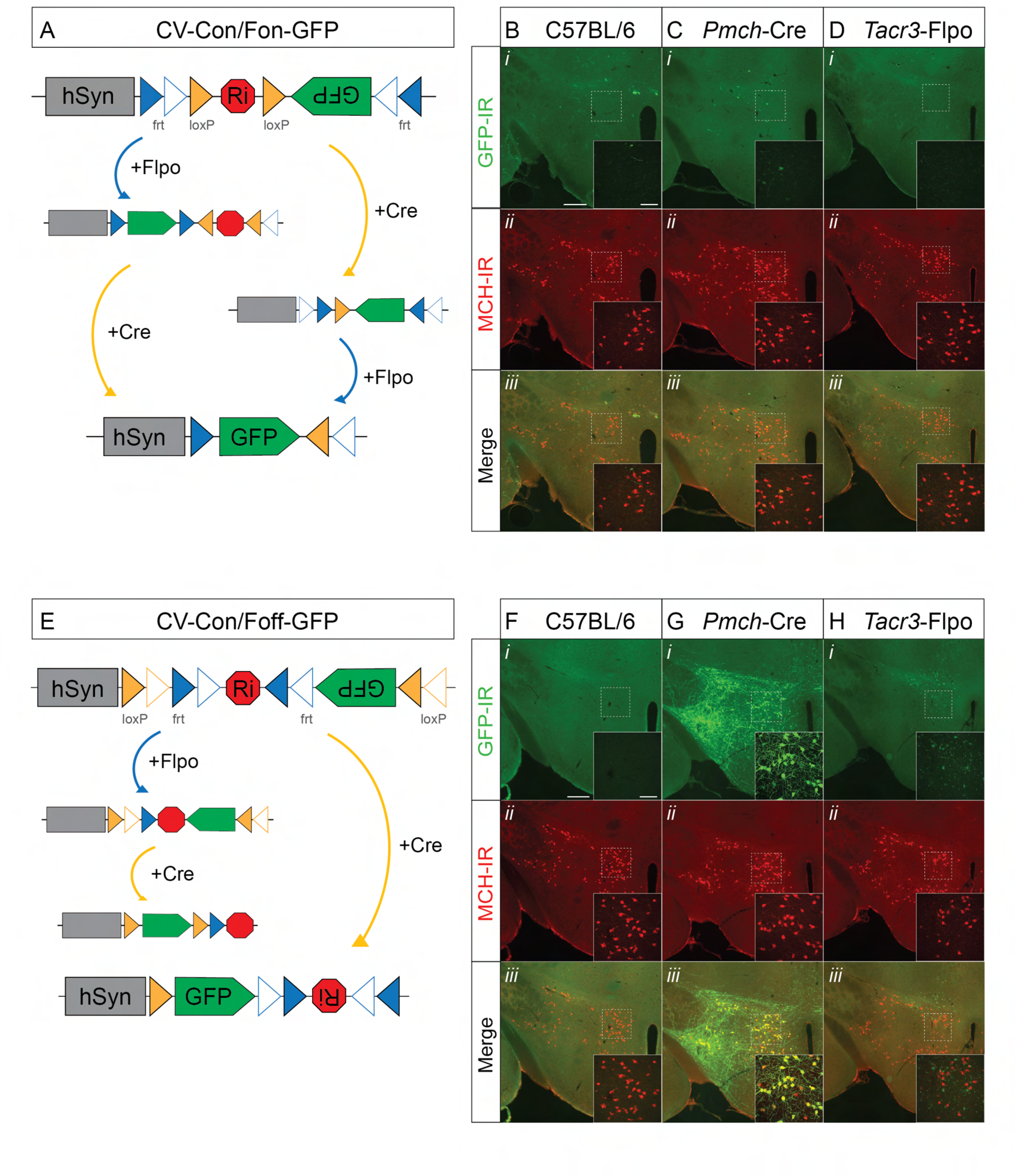
Validation of Conditional Viral Expression by Ribozyme Guided Degradation (ConVERGD) viruses. **A)** Schematic representing AAV2/8-hSyn-CV-Con/Fon-GFP viral construct and recombination in the presence no recombinase, Flpo recombinase only, Cre recombinase only, and Flpo AND Cre recombinase conditions. In the presence of no Flpo or Cre recombinase, the ribozyme and inverted GFP cassette will prevent GFP expression to occur. In the presence of Flpo recombinase ONLY, Flpo recombination will invert the GFP cassette, but GFP expression will be inhibited by the presence of the ribozyme (red stop symbol). In the presence of Cre recombinase ONLY, Cre recombination will excise the ribozyme, but the GFP cassette will still be inverted and no GFP expression will occur. In the presence of Flpo AND Cre recombinase, the GFP cassette will be inverted through Flpo mediated recombination, and the ribozyme will be excised through Cre mediated recombination, allowing GFP expression to occur. Viral construct schematics adapted from Hughes et al., 2024. **B-D)** AAV2/8-hSyn-CV-Con/Fon-GFP viral injection in C57BL/6 **(B)**, *Pmch*-Cre **(C)**, and *Tacr3*-Flpo **(D)** control animals. Representative epifluorescent photomicrograph (10X) of GFP-IR (green; ***i***), MCH-IR (red; ***ii***), and merge (***iii***) of LHA from injected control animals. Insets represent confocal photomicrographs (40X) from outlined area in epifluorescent image. **E)** Schematic representing AAV2/8-hSyn-CV-Con/Foff-GFP viral construct in the presence of no recombinase, Flpo recombinase only, Cre recombinase only, and Flpo AND Cre recombinase conditions. In the presence of no Flpo and Cre recombinase, the ribozyme and inverted GFP cassette will prevent GFP expression to occur. In the presence of Flpo recombinase ONLY, the ribozyme (stop symbol) will be inverted, and as the GFP cassette remains inverted, the expression of GFP will not occur. In the presence of Cre AND Flpo recombinase, the GFP cassette will be inverted, but the ribozyme will be un-inverted, thus preventing GFP expression to occur. In the presence of Cre recombinase and NOT Flpo recombinase, Cre-mediated recombination will invert both GFP and ribozyme cassettes, and GFP expression will occur. Viral construct schematic adapted from Hughes et al., 2024. **F-G)** AAV2/8-hSyn-CV-Con/Foff-GFP viral injection in C57BL/6 **(F)**, *Pmch*-Cre **(G)**, and *Tacr3*-Flpo **(H)** control animals. Representative epifluorescent photomicrograph (10X) of GFP-IR (green; ***i***), MCH-IR (red; ***ii***), and merge (***iii***) of LHA from injected control animals. Insets represent confocal photomicrographs (40X) from outlined area in epifluorescent image. 10X scale bars: 300 µM. 40X scale bars: 100 µM.

### Validation of injection site and viral expression in MCH neuron subpopulations

We next assessed viral specificity at the bilateral injection sites across the three experimental conditions in male and female mice as described in Fig. 7.In *Pmch*-Cre animals injected with DIO-EYFP, we quantified specificity by immunostaining for MCH and GFP in order to enhance DIO-EYFP viral expression **(Fig. 9A-B).** Nearly all (99.4%) DIO-EYFP cells colocalized with MCH-IR throughout the LHA (2094 EYFP cells; N=3) **(Fig. 9C)**, and approximately 84.6% of MCH-IR neurons expressed DIO-EYFP (2462 MCH-IR cells; N=3) (**Fig. 9D**). This high injection penetrance is consistent with previous reports of low non-specific Cre expression in the *Pmch*-Cre line (Beekly et al., 2020; Miller et al., 2024). As a marker for Type I MCH neurons, we next quantified the proportion of DIO-EYFP cells that co-expressed CART-IR **(Fig. 9E-F).** We found that 47.3% of DIO-EYFP neurons expressed CART-IR (1751 DIO-EYFP cells; N=3) **(Fig. 9G)**. Additionally, we found high DIO-EYFP penetrance of the total CART-IR population, with 81.1% of total CART-IR cells expressing DIO-EYFP (1022 total CART-IR cells; N=3) **(Fig. 9H).**

**Figure 9:**
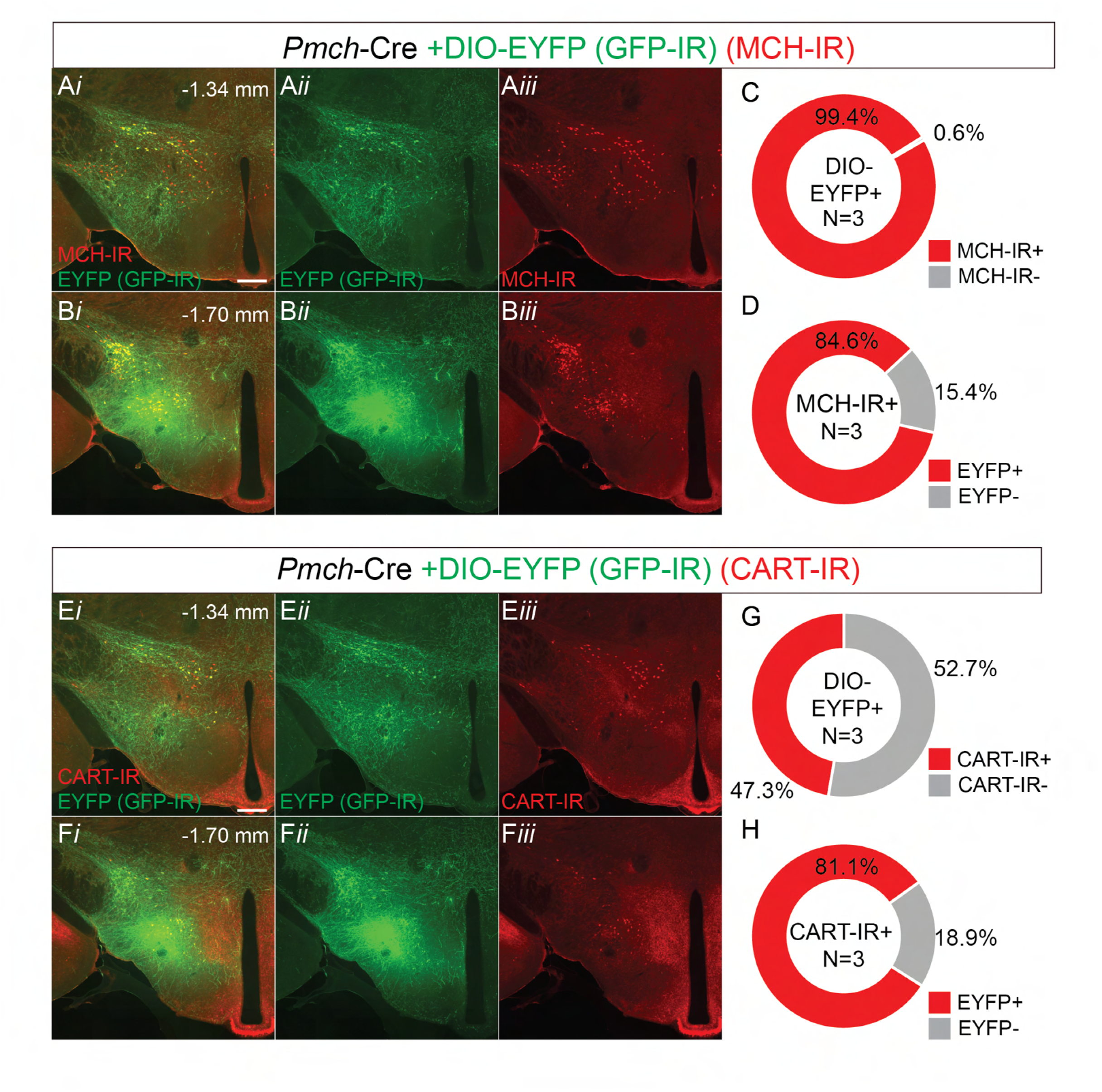
DIO-EYFP expression is restricted to MCH-IR neurons in the LHA of *Pmch*-Cre mice. A-B) Representative epifluorescent (10X) photomicrograph of AAV2-Ef1a-DIO-EYFP (+GFP-IR) injection site, counterstained against MCH, in the *Pmch*-Cre animal in anterior (−1.34 mm from bregma, **A**) and posterior (−1.70 mm from bregma, **B**) sections of the LHA. Panels show merged (***i***), DIO-EYFP only (+GFP-IR, green, ***ii***), and MCH-IR only (red, ***iii***) photomicrographs. **C)** Donut plot representing DIO-EYFP specificity to MCH-IR neurons (2094 total EYFP cells; N=3). **D)** Donut plot representing DIO-EYFP penetrance of total MCH-IR neurons (2462 total MCH-IR cells; N=3). **E-F)** Merged representative epifluorescent (10X) photomicrograph of AAV2-Ef1a-DIO-EYFP (+GFP-IR) injection site, counterstained against CART, in the *Pmch*-Cre animal in anterior (−1.34 mm from bregma, **E**) and posterior (−1.70 mm from bregma, **F**) sections of the LHA. Panels show merged (***i***), DIO-EYFP only (+GFP-IR, green; ***ii***), and CART-IR only (red; ***iii***) photomicrographs. **G)** Donut plot representing EYFP specificity to CART-IR neurons (1751 total DIO-EYFP cells; N=3). **H)** Donut plot representing EYFP penetrance of total CART-IR neurons (1022 total CART-IR cells; N=3). 10X scale bars: 300 µM. Sections in A and E, and B and F are adjacent and taken from the same animal to allow for direct comparison of GFP-IR with MCH-IR and CART-IR.

Next, we assessed intersectional targeting of Type I MCH neurons by injecting *Pmch*-Cre; *Tacr3*-Flpo animals with CV-Con/Fon-GFP virus (**Fig 10**). CV-Con/Fon-GFP+ neurons were more densely distributed in anterior than posterior sections of the LHA **(Fig. 10A-B, E-F)** and is consistent with previous reports of Type I MCH neuron distribution. We found that 90.9% of CV-Con/Fon-GFP expressing neurons colocalized with MCH-IR throughout the LHA (n=1145 CV-Con/Fon-GFP cells; N=3) **(Fig. 10C**). Similarly, 84.9% of CV-Con/Fon-GFP cells co-expressed CART (n=835 CV-Con/Fon-GFP cells; N=3), indicating high specificity to Type I MCH neurons **(Fig. 10G).** Furthermore, the virus exhibited high penetrance, with 59.8% of total MCH-IR and 79.4% of total CART-IR+ neurons co-expressing CV-Con/Fon-GFP (n= 1742 MCH-IR cells; n=893 CART-IR cells; N=3) (**Fig. 10D**, H).

**Figure 10:**
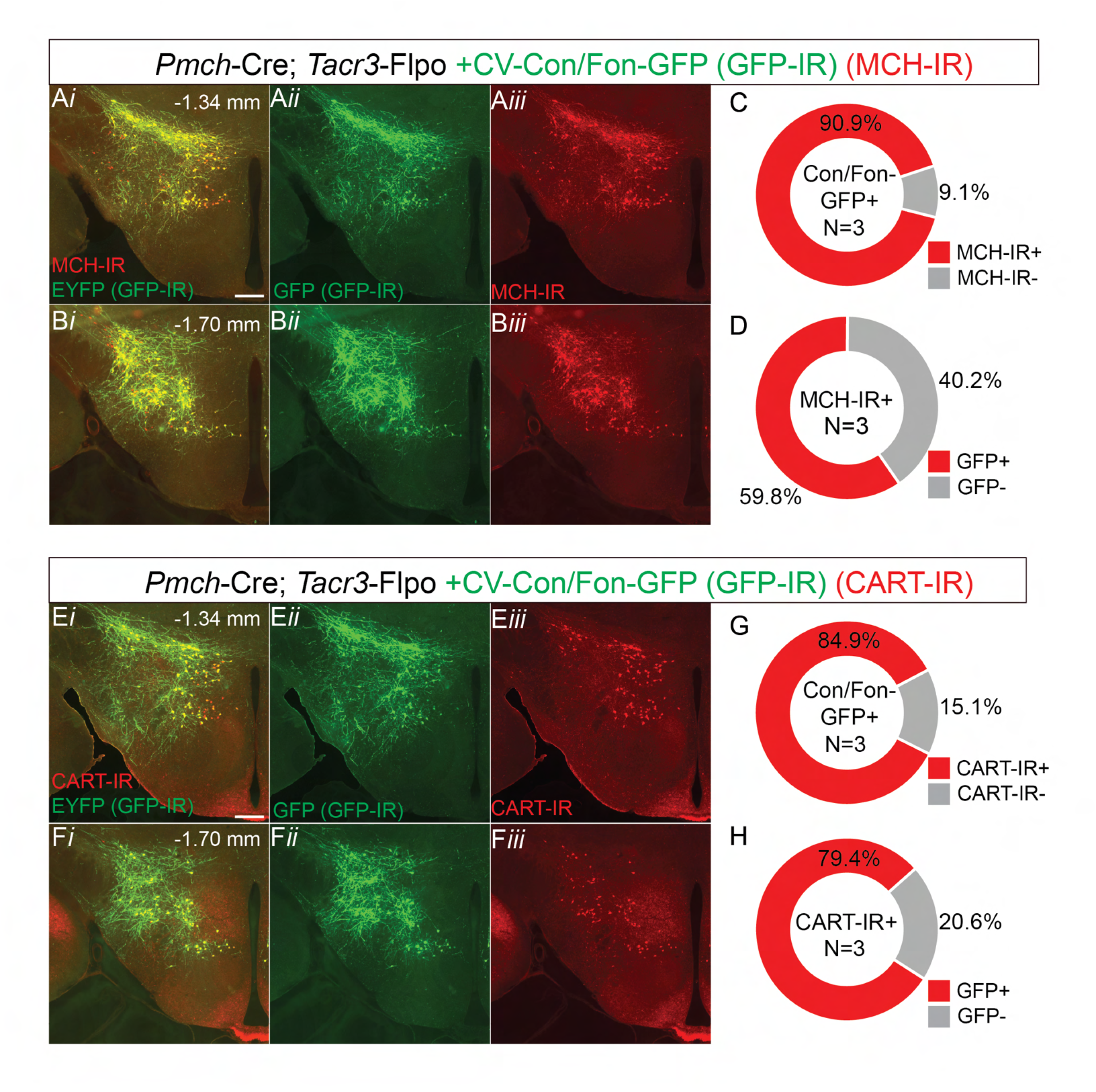
CV-Con/Fon-GFP selectively labels Type I MCH neurons in the LHA of dual recombinase animals. A-B) Representative epifluorescent (10X) photomicrograph of AAV2/8-hSyn-CV(Con/Fon)-GFP (+GFP-IR) injection site, counterstained against MCH, in the *Pmch*-Cre; *Tacr3*-Flpo animal in anterior (−1.34 mm from bregma, **A**) and posterior (−1.70 mm from bregma, **B**) sections of the LHA. Panels show merged (***i***), CV-Con/Fon-GFP only (+GFP-IR, green; ***ii***), and MCH-IR only (red; ***iii***) photomicrographs. **C)** Donut plot representing CV-Con/Fon-GFP specificity to MCH-IR neurons (1145 total CV-Con/Fon-GFP cells; N=3). **D)** Donut plot representing CV-Con/Fon-GFP penetrance of total MCH-IR neurons (1742 total MCH-IR cells; N=3). **E-F)** Merged representative epifluorescent (10X) photomicrograph of AAV28-hSyn-CV(Con/Fon)-GFP (+GFP-IR) injection site, counterstained against CART, in the *Pmch*-Cre; *Tacr3*-Flpo animal in anterior (−1.34 mm from bregma, **E**) and posterior (−1.70 mm from bregma, **F**) sections of the LHA. Panels show merged (***i***), CV-Con/Fon-GFP only (+GFP-IR, green;***ii***), and CART-IR only (red;***iii***) photomicrographs. **G)** Donut plot representing CV-Con/Fon-GFP specificity to CART-IR neurons (835 total CV-Con/Fon-GFP cells; N=3). **H)** Donut plot representing CV-Con/Fon-GFP penetrance of total CART-IR neurons (893 total CART-IR cells; N=3). 10X scale bars: 300 µM. Sections in A and E are adjacent and taken from the same animal to allow for direct comparison of GFP-IR with MCH-IR and CART-IR.

Finally, we assessed intersectional targeting of Type II MCH neurons by injecting *Pmch*-Cre;*Tacr3*-Flpo animals with CV-Con/Foff-GFP virus (**Fig. 11**). Qualitatively, CV-Con/Foff-GFP+ neurons were denser in posterior and lateral regions of the LHA, consistent with previous reports of Type II MCH neuron distribution (**Fig 11A-B, E-F**). A majority (89.5%) of CV-Con/Foff-GFP+ neurons colocalized with MCH-IR+ neurons and labeled 37.9% of MCH-IR neurons (1067 CV-Con/Foff-GFP cells; n=2521 MCH-IR cells; N=3) **(Fig. 11C-D).** Furthermore, we found that 73.7% of CV-Con/Foff-GFP+ neurons did not colocalize with CART-IR, confirming preferential targeting to Type II MCH neurons (n=1133 CART-IR cells; N=3) **(Fig. 11G).** Additionally, only 21.8% CART-IR cells expressed CV-Con/Foff-GFP (1364 total CART-IR cells; N=3), consistent with our expectation that CART expression across MCH subpopulations is a spectrum rather than purely binary and that a subset of Type II MCH neurons express CART-IR.

**Figure 11:**
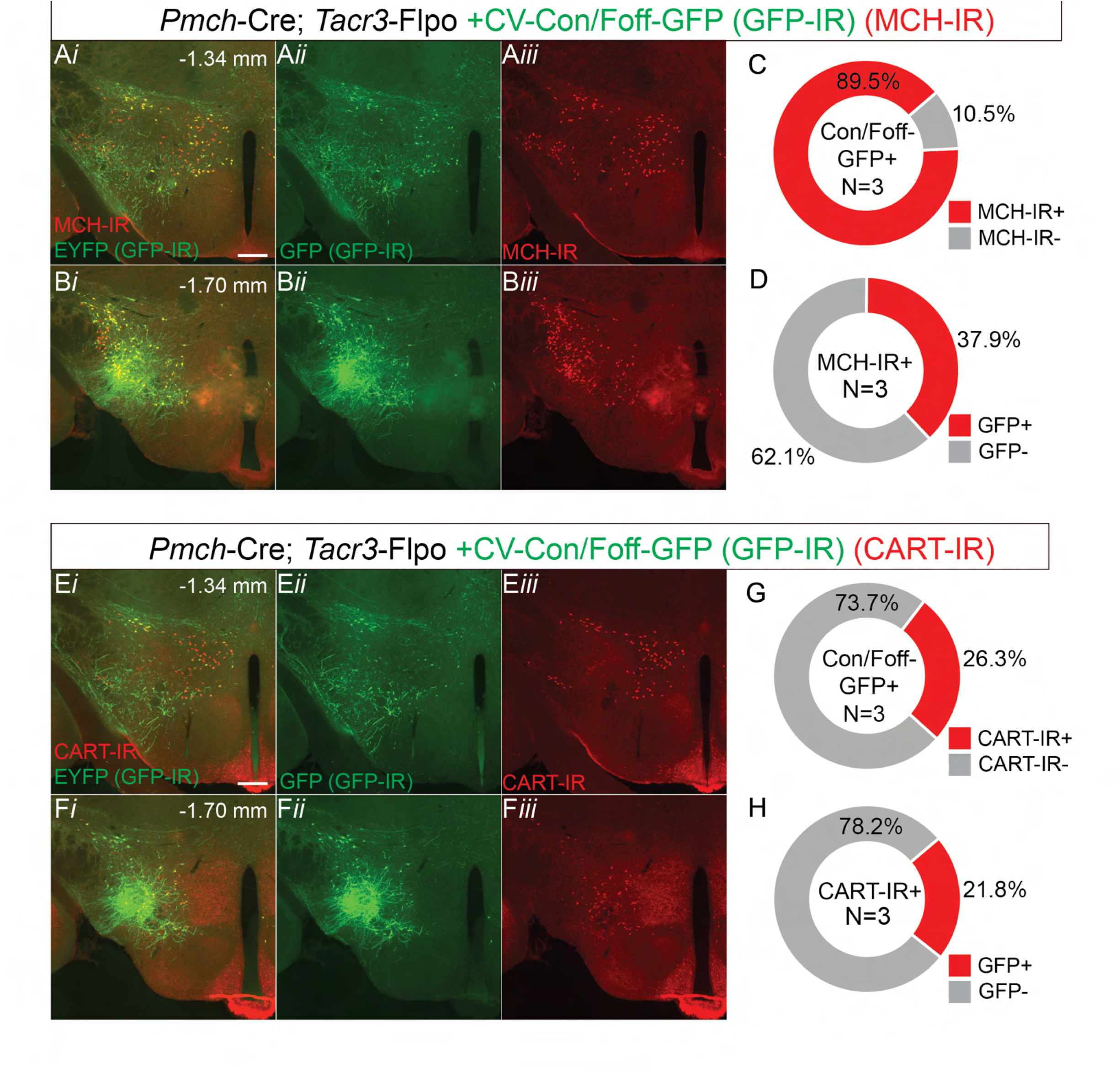
CV-Con/Foff-GFP selectively labels Type II MCH neurons in the LHA of dual recombinase animals. A-B) Representative epifluorescent (10X) photomicrograph of AAV28-hSyn-CV(Con/Foff)-GFP (+GFP-IR) injection site, counterstained against MCH, in the *Pmch*-Cre; *Tacr3*-Flpo animal in anterior (−1.34 mm from bregma, **A**) and posterior (−1.70 mm from bregma, **B**) sections of the LHA. Panels show merged (***i***), CV-Con/Foff-GFP only (+GFP-IR, green; ***ii***), and MCH-IR only (red; ***iii***) photomicrographs. **C)** Donut plot representing CV-Con/Foff-GFP specificity to MCH-IR neurons (1067 total CV-Con/Foff-GFP cells; N=3). **D)** Donut plot representing CV-Con/Foff-GFP penetrance of total MCH-IR neurons (2521 total MCH-IR cells; N=3). **E-F)** Merged representative epifluorescent (10X) photomicrograph of AAV28-hSyn-CV(Con/Foff)-GFP (+GFP-IR) injection site, counterstained against CART, in the *Pmch*-Cre; *Tacr3*-Flpo animal in anterior (−1.34 mm from bregma, **E**) and posterior (−1.70 mm from bregma, **F**) sections of the LHA. Panels show merged (***i***), CV-Con/Foff-GFP only (+GFP-IR, green; ***ii***), and CART-IR only (red; ***iii***) photomicrographs. **G)** Donut plot representing CV-Con/Foff-GFP specificity to CART-IR neurons (1133 total CV-Con/Foff-GFP cells; N=3). **H)** Donut plot representing CV-Con/Foff-GFP penetrance of total CART-IR neurons (1364 total CART-IR cells; N=3). 10X scale bars: 300 µM. Sections in A and E, and B and F are adjacent and taken from the same animal to allow for direct comparison of GFP-IR with MCH-IR and CART-IR.

Having confirmed viral specificity and differential spatial distribution of intersectionally targeted neurons across all three conditions, we next compared the anterograde projections patterns of total MCH, Type I, and Type II MCH neurons throughout the mouse brain.

### Viral anterograde projection comparison of Type I and Type II neurons reveal diverging and converging circuits

We qualitatively assessed and compared the density of anterograde labeled fibers across matched coronal sections, spanning key regions across the antero-posterior extent of the brain in DIO-EYFP (total MCH), CV-Con/Fon-GFP (Type I MCH), and CV-Con/Foff-GFP (Type II MCH) injected brains (N=3 per condition), while referencing Allen Adult Mouse Brain Atlas (atlas.brain-map.org) for neuroanatomical registration **(Figs. 12-15).** Virally mediated EYFP or GFP fluorescence was amplified with GFP-IR and computationally cleared to enhance the visualization of fine axonal processes. To characterize and compare fiber density across conditions, fiber density was qualitatively scored from a range of - (little to no coverage), + (sparse), ++ (moderate), +++ (dense), and ++++ (very dense) as shown in **Figure 16**. Finally, we did not observe obvious sex differences in the pattern or density of anatomical projections across the three conditions, consist with previous work showing that MCH subpopulation cell body count does not differ between male and female mice (Miller et al., 2024).

**Figure 12:**
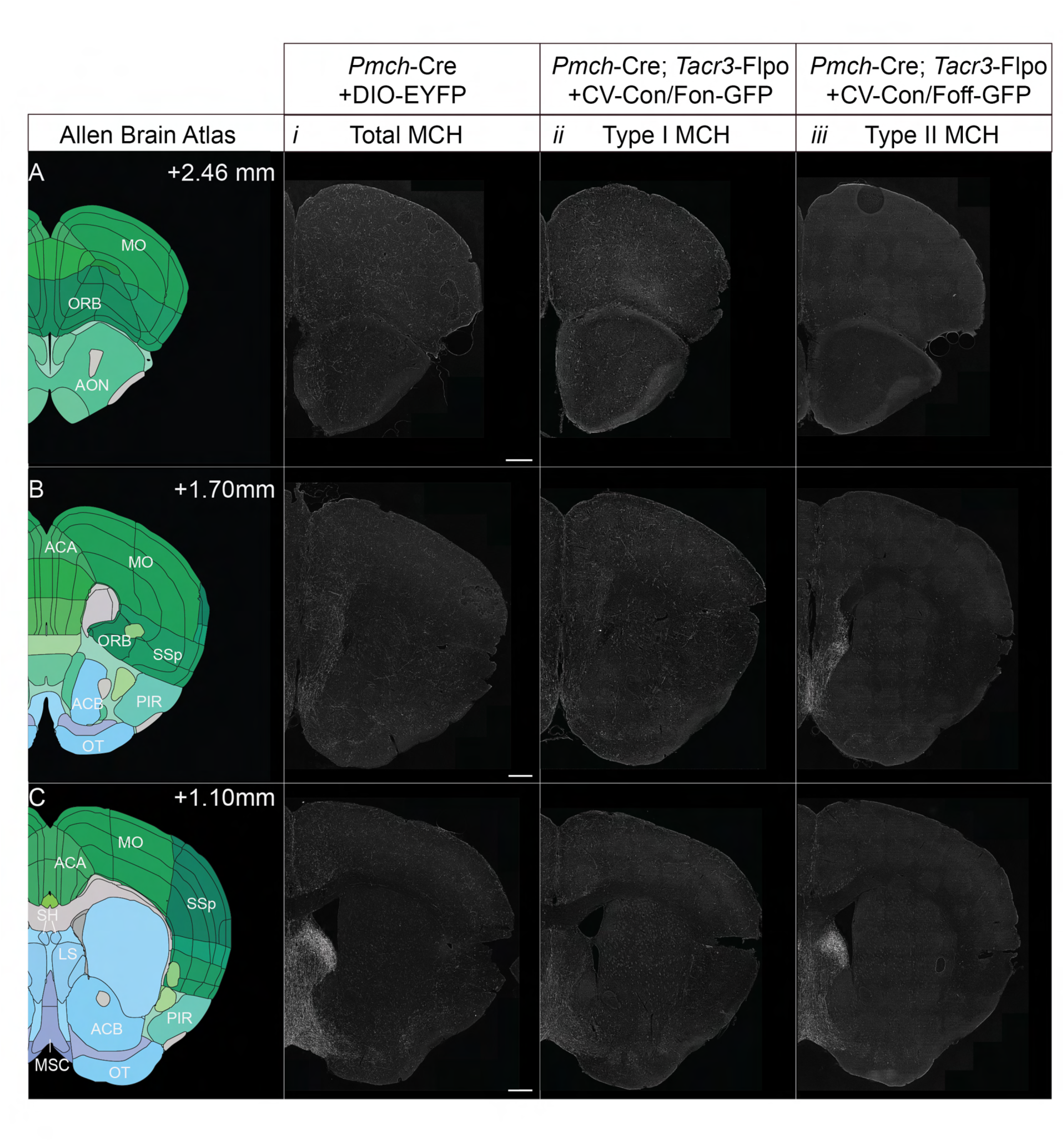
Anterograde projections of total, Type I, and Type II MCH neurons in the forebrain. (A-C) Representative coronal sections at three anatomical levels (distance from bregma: +2.46, +1.70, and +1.10 mm, respectively). Each row represents corresponding level taken from the Allen Adult Mouse Brain Atlas (//atlas.brain-map.org/; left), followed by representative 20X Thunder cleared fluorescent images taken from three experimental conditions: *Pmch*-Cre+DIO-EYFP (female, total MCH projections; ***i***), *Pmch*-Cre; *Tacr3*-Flpo +CV-Con/Fon-GFP (male, Type I MCH projections; ***ii***), and *Pmch*-Cre; *Tacr3*-Flpo +CV-Con/Foff-GFP (female, Type II MCH projections; ***iii***). Figures 12-15 show representative sections taken from the same animal per experimental condition to enable direct within animal comparison. **Abbreviations:** ACB: nucleus accumbens; ACA: anterior cingulate area; AON: anterior olfactory nucleus; IL: infralimbic area; LS: lateral septum; MS: medial septum; MO: somatomotor areas; NDB: nucleus of the diagonal band; ORB: orbital area; PIR: piriform area; PL: prelimbic area. Scale bars represent 500 µM.

**Figure 13:**
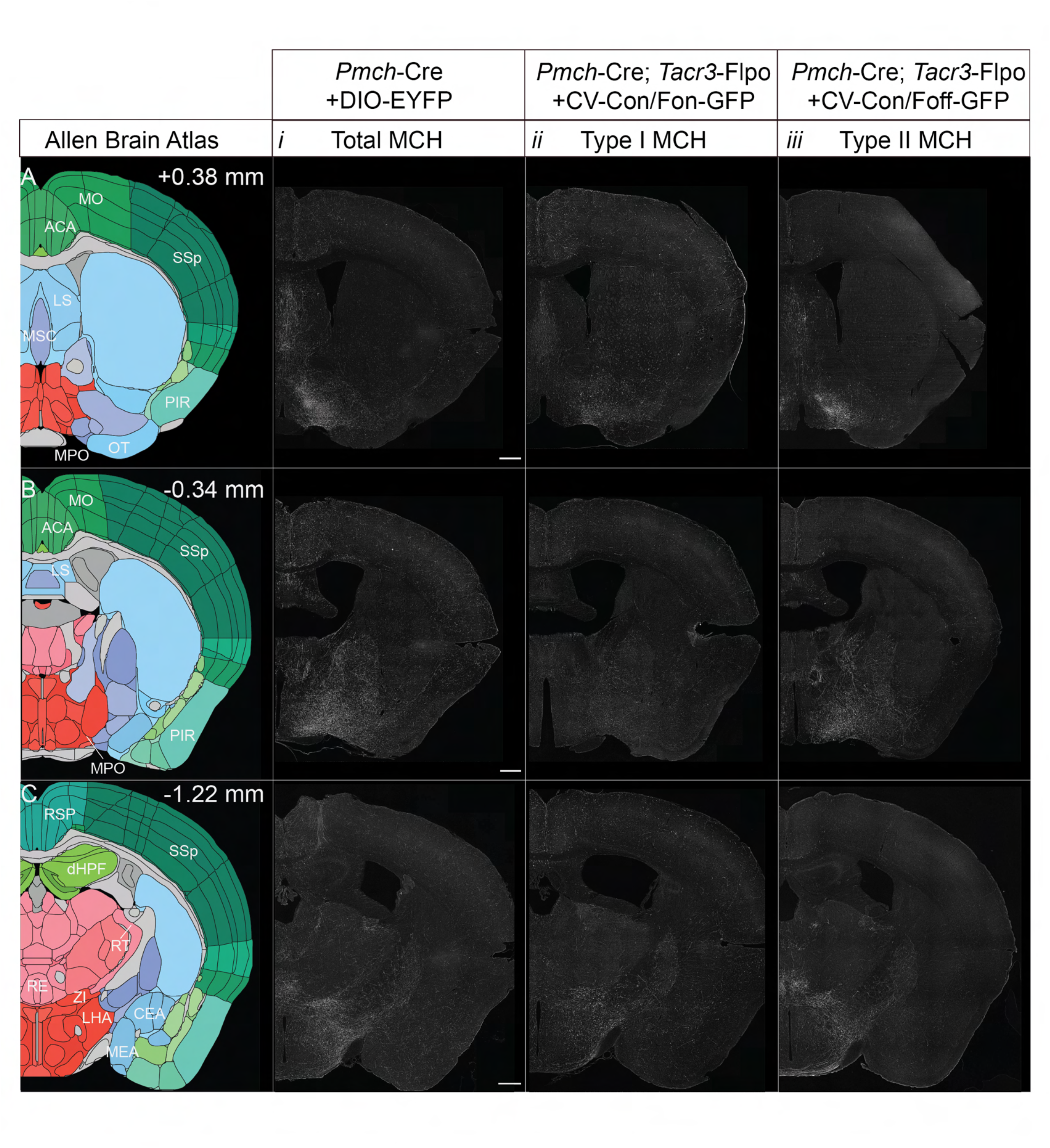
Anterograde projections of total, Type I, and Type II MCH neurons in the basal forebrain and hypothalamus. (A-C) Representative coronal sections at three anatomical levels (distance from bregma: +0.38, -0.34, and -1.22 mm, respectively). Each row represents corresponding level taken from the Allen Adult Mouse Brain Atlas (//atlas.brain-map.org/; left), followed by representative 20X Thunder cleared fluorescent images taken from three experimental conditions: *Pmch*-Cre+DIO-EYFP (female, total MCH projections; ***i***), *Pmch*-Cre; *Tacr3*-Flpo +CV-Con/Fon-GFP (male, Type I MCH projections; ***ii***), and *Pmch*-Cre; *Tacr3*-Flpo +CV-Con/Foff-GFP (female, Type II MCH projections; ***iii***). Figures 12-15 show representative sections taken from the same animal per experimental condition to enable direct within animal comparison. **Abbreviations:** ACA: anterior cingulate area; AVPV: anteroventral periventricular nucleus; dHPF: dorsal hippocampal formation; GP: globus pallidus; GPe: external segment of the globus pallidus; ILM: intralaminar thalamus; LHA: lateral hypothalamic area; MOp: primary motor area; MPO: medial preoptic area; PIR: piriform area; RE: nucleus reuniens; SF: septofimbrial nucleus; SI: substantia innominata; SSp: primary somatosensory area. Scale bars represent 500 µM.

**Figure 14:**
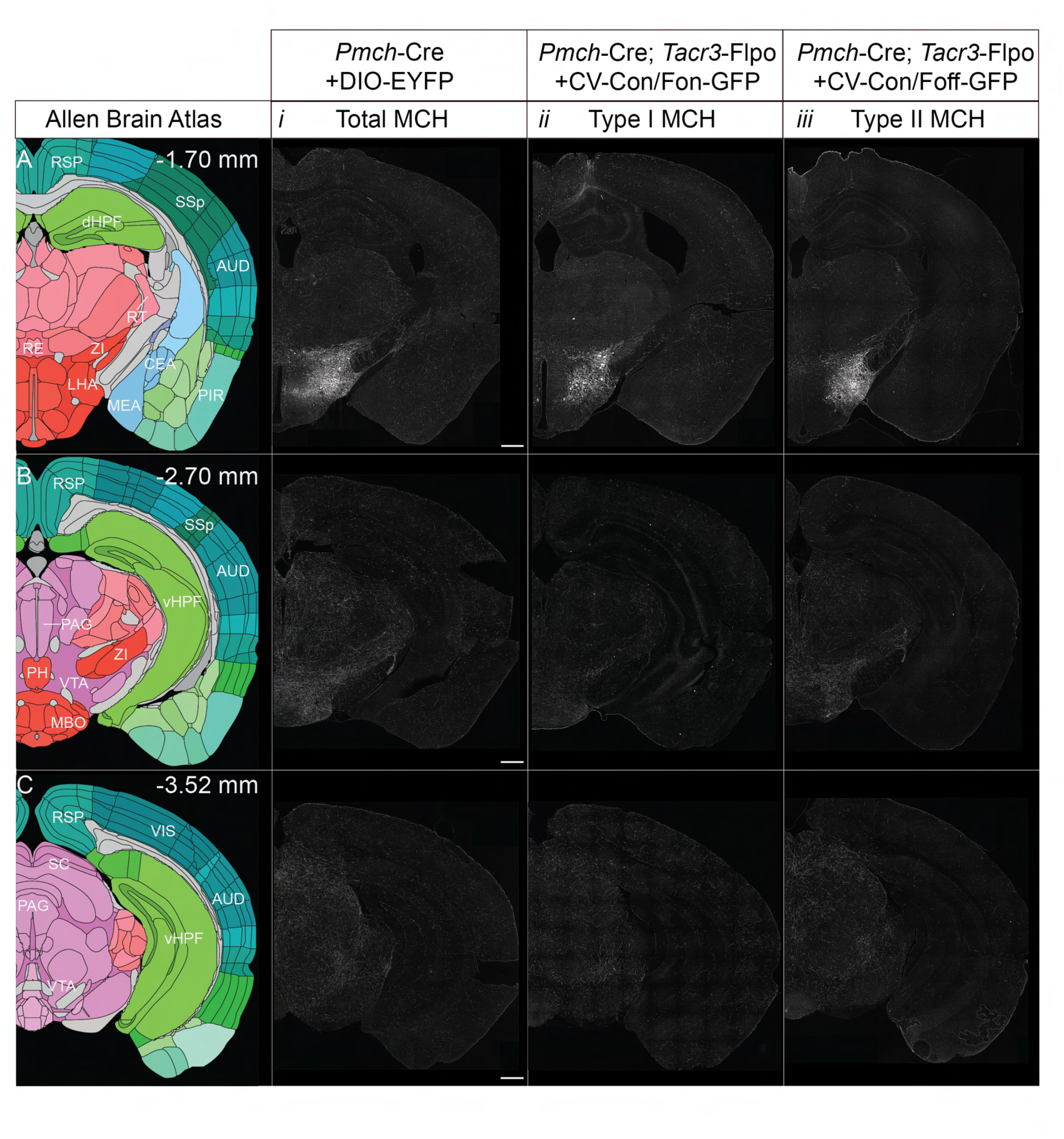
Anterograde projections of total, Type I, and Type II MCH neurons in the hypothalamus and midbrain. (A-C) Representative coronal sections at three anatomical levels (distance from bregma: -1.70, -2.70, and -3.52 mm, respectively). Each row represents corresponding level taken from the Allen Adult Mouse Brain Atlas (//atlas.brain-map.org/; left), followed by representative 20X Thunder cleared fluorescent images taken from three experimental conditions: *Pmch*-Cre+DIO-EYFP (female, total MCH projections; ***i***), *Pmch*-Cre; *Tacr3*-Flpo +CV-Con/Fon-GFP (male, Type I MCH projections; ***ii***), and *Pmch*-Cre; *Tacr3*-Flpo +CV-Con/Foff-GFP (female,Type II MCH projections; ***iii***). Figures 12-15 show representative sections taken from the same animal per experimental condition to enable direct within animal comparison, except for (***Cii***) representative section is shown from a separate mouse (female) from (*Aii)* and (*Bii*). **Abbreviations:** dHPF: dorsal hippocampal formation; LHA: lateral hypothalamic area; LM: lateral mammillary nucleus; MRN: midbrain reticular nucleus; MM: medial mammillary nucleus; PAG: periaqueductal grey; RT: reticular nucleus of the thalamus; RSP: retrosplenial area; SC: superior colliculus; STN: subthalamic nucleus; vHPF: ventral hippocampal formation; VTA: ventral tegmental area; ZI: zona incerta. Scale bars represent 500 µM.

**Figure 15:**
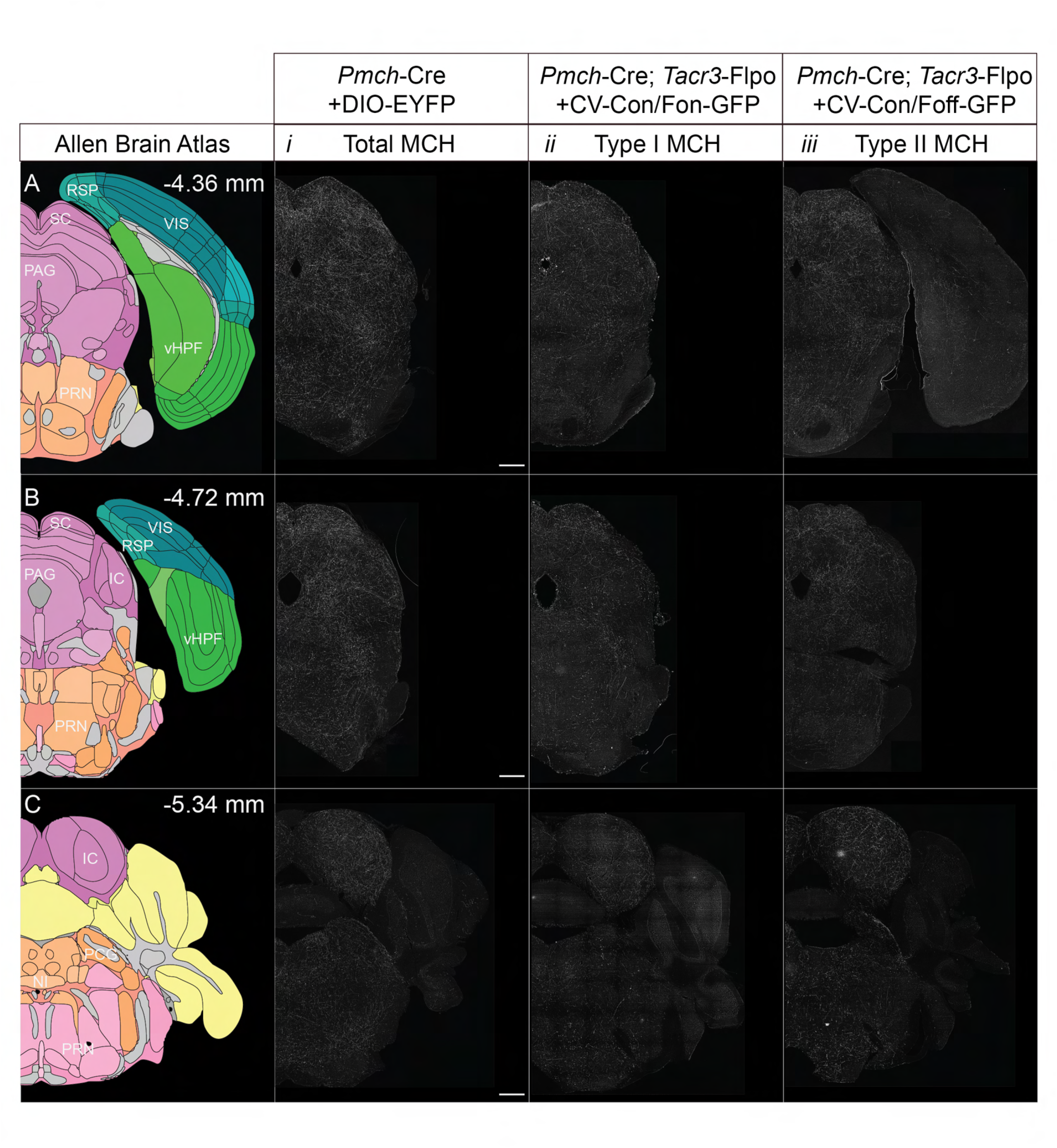
Anterograde projections of total, Type I, and Type II MCH neurons in the hindbrain. (A-C) Representative coronal sections at three anatomical levels (distance from bregma: -4.36, -4.72, and –5.34 mm, respectively). Each row represents corresponding level taken from the Allen Adult Mouse Brain Atlas (//atlas.brain-map.org/; left), followed by representative 20X Thunder cleared fluorescent images taken from three experimental conditions: *Pmch*-Cre+DIO-EYFP (female, total MCH projections; ***i***), *Pmch*-Cre; *Tacr3*-Flpo +CV-Con/Fon-GFP (male, Type I MCH projections; ***ii***), and *Pmch*-Cre; *Tacr3*-Flpo +CV-Con/Foff-GFP (female, Type II MCH projections; ***iii*** Figures 12-15 show representative sections taken from the same animal per experimental condition to enable direct within animal comparison, except for (***Cii***) representative section is shown from a separate mouse (female) from (*Aii)* and (*Bii*). **Abbreviations:** IC: inferior colliculus; MRN: midbrain reticular nucleus; NI: nucleus incertus; PAG: periaqueductal grey; PBN: parabrachial nucleus; PCG: pontine central grey; PG: pontine grey; PRN: pontine reticular nucleus; SC: superior colliculus ; TRN: thalamic reticular nucleus. Scale bars represent 500 µM.

**Figure 16:**
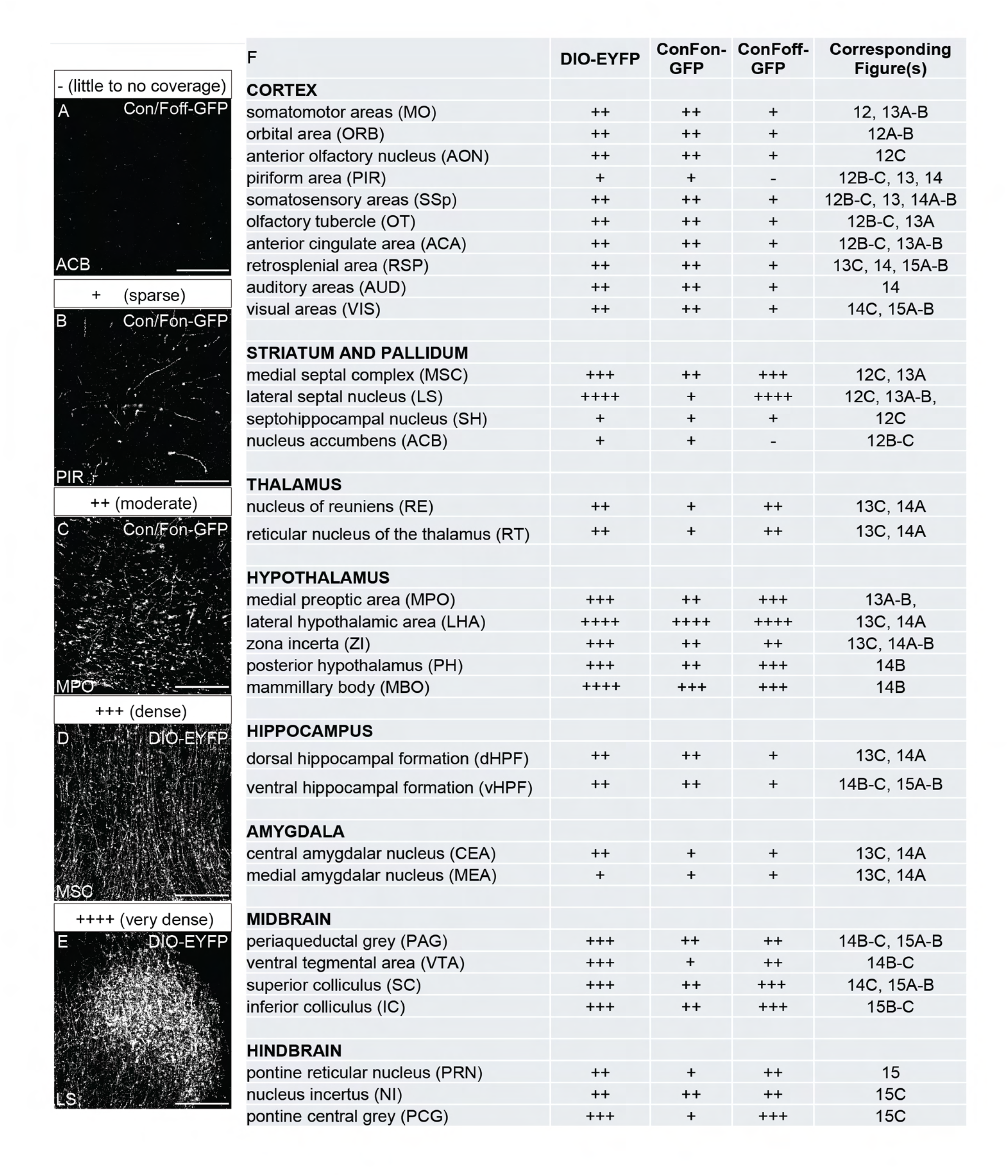
Summary of fiber analysis of total MCH, Type I MCH, and Type II MCH neurons (A-E) Representative 20X epifluorescent photomicrographs, illustrating each level of fiber density scoring: - (little to no coverage) exemplified by CV-Con/Foff-GFP fibers in the nucleus accumbens (ACB); + (sparse) exemplified by CV-Con/Fon-GFP fibers in the piriform area (PIR); ++ (moderate), exemplified by CV-Con/Fon-GFP fibers in the medial preoptic area (MPO); +++ (dense), exemplified by DIO-EYFP fibers in the medial septal complex (MSC); ++++ (very dense), exemplified by DIO-EYFP fibers in the lateral septum (LS). **(F)** Qualitative fiber density score from total MCH (*Pmch*-Cre+DIO-EYFP), Type I MCH (*Pmch*-Cre; *Tacr3*-Flpo +CV-Con/Fon-GFP) and Type II MCH (*Pmch*-Cre; *Tacr3*-Flpo +CV-Con/Foff-GFP) projections organized by subregions and observed across brains (N=3 per condition). Fiber density was assessed on a five-point scale: - (little to no coverage), + (sparse), ++ (moderate), +++ (dense), and ++++ (very dense). Brain region abbreviations follow the Allen Adult Mouse Brain Atlas abbreviations. Scale bars represent 250 µM.

In anterior sections **(Fig. 12),** moderate total and Type I MCH fiber labeling was observed in the somatomotor (MO) and orbital areas (ORB) of the cortex, while Type II fiber signal was sparse in these regions. Similarly, other cortical subregions like the anterior olfactory nucleus (AON), somatosensory area (SSp), and olfactory tubercle (OT) also demonstrated similar total and Type I MCH fibers, with sparse Type II MCH fibers. The piriform area (PIR) of the cortex demonstrated sparse total and Type I MCH fiber signal but very little to no Type II MCH fibers. Overall, these anterior sections demonstrate preferential cortical innervation by Type I MCH neurons.

Within septal and pallidal structures (**Fig. 12B-C, 13A-B)**, we found moderately dense to very dense total MCH fibers in the medial septal complex (MSC), which comprised the medial septum and nucleus of the diagonal band, and the lateral septal nucleus (LS). These areas contained the strongest signals from Type II MCH fibers and only moderate or sparse signals from Type I MCH fibers. By contrast, the septohippocampal nucleus (SH) and nucleus accumbens (ACB) contained sparse total MCH and Type I MCH signal or sparse to no Type II MCH signal.

Next, at the level of anterior hypothalamic and thalamic structures **(Figs. 13, 14A-B)**, the medial preoptic area (MPO) contained dense total MCH and Type II MCH fibers but moderate Type I MCH fibers. The LHA and zona incerta (ZI) demonstrated very dense fiber labeling across all three conditions and is consistent with local axonal processes from MCH cell bodies at the virus injection site. Other hypothalamic structures, including the posterior hypothalamus (PH) and mammillary body (MBO), contained dense total MCH and Type II MCH fibers but moderate Type I MCH fibers (Figs. 14A-B). Interestingly, fiber distribution within the MBO appeared denser laterally compared to medially. Total MCH fibers in the thalamus were relatively sparse compared to the hypothalamus, and only the nucleus reuniens (RE) and reticular nucleus of the thalamus (RT) exhibited notable fibers from MCH neurons. The RE and RT demonstrated moderate total MCH and Type II labeling but sparse Type I signal. Together, thalamic targets, particularly in the RE, reflected preferential innervation from Type II MCH neurons compared with Type I.

Both the dorsal and ventral hippocampal formation **(Figs. 13C,14,15A-B)** demonstrated moderate to sparse total MCH fiber signals. This signal was largely dominated by Type I MCH fibers, which contributed moderate fiber signals, while Type II MCH signals were sparse in both subregions. Additionally, the central and medial amygdalar nuclei **(Figs. 13C,14A)** exhibited sparse to moderate total MCH signal, with sparse Type I and Type II MCH fiber presence.

Cortical regions in posterior sections **(Figs. 13-15)** continued to demonstrate a Type I MCH dominant pattern as observed in anterior sections. The retrosplenial area (RSP), auditory area (AUD), and visual area (VIS) of the cortex contained moderate total MCH and sparse Type I labeling with sparse Type II signal.

Midbrain structures **(Figs. 14B-C, 15)**, such as the periaqueductal grey (PAG), ventral tegmental area (VTA), superior colliculus (SC), and inferior colliculus (IC) all exhibited dense total MCH innervation. The PAG contained similar moderate innervation of Type I and Type II MCH neurons. while the VTA contained sparse Type I MCH signal but moderate Type II MCH signal. By contrast, the SC and IC contained moderate Type I MCH signal, yet dense Type II MCH signal.

Brainstem nuclei **(Fig. 15)** exhibited moderate to dense total MCH fibers. The pontine reticular nucleus (PRN) and pontine central grey (PCG) were preferentially innervated by Type II MCH neurons, with moderate to dense Type II fiber labeling and sparse Type I fiber labeling. However, the nucleus incertus (NI) displayed equally moderate Type I and Type II fiber innervation.

Overall, these findings, summarized in **Figure 16**, demonstrate that Type I and Type II MCH neurons extended largely overlapping projection targets, yet with evidence of preferential innervation suggestive of distinct subcircuits in both male and female mice.

### High-spatial resolution fiber mapping of total MCH, Type I MCH, and Type II MCH fibers in the septal complex and cortex

We next performed high resolution fiber mapping in the septal complex and adjacent cortical subregions to directly compare the projection patterns between MCH subpopulations and determine if differences in their fiber patterns persist across sex. As previously described (Bono et al., 2025; Miller et al., 2024; Negishi et al., 2020; Payant et al., 2024), we generated representative maps of fiber distribution in the cortex and septal complex corresponding to anterior, middle, and posterior, levels (distance from bregma: +1.245, +0.745, and +0.145 mm, respectively) in male (**Fig. 17**) and female mice (**Fig. 18**). As we mapped only the medial prefrontal cortex and septal complex, we omitted fibers in the taenia tecta (TTd), caudoputamen (CP), nucleus accumbens (ACB), substantia innominata (SI), olfactory tubercle (OT), and bed nucleus of the stria terminalis (BST), and hypothalamic areas from these maps; their absence reflects an intentional omission rather than a biological phenomenon.

**Figure 17:**
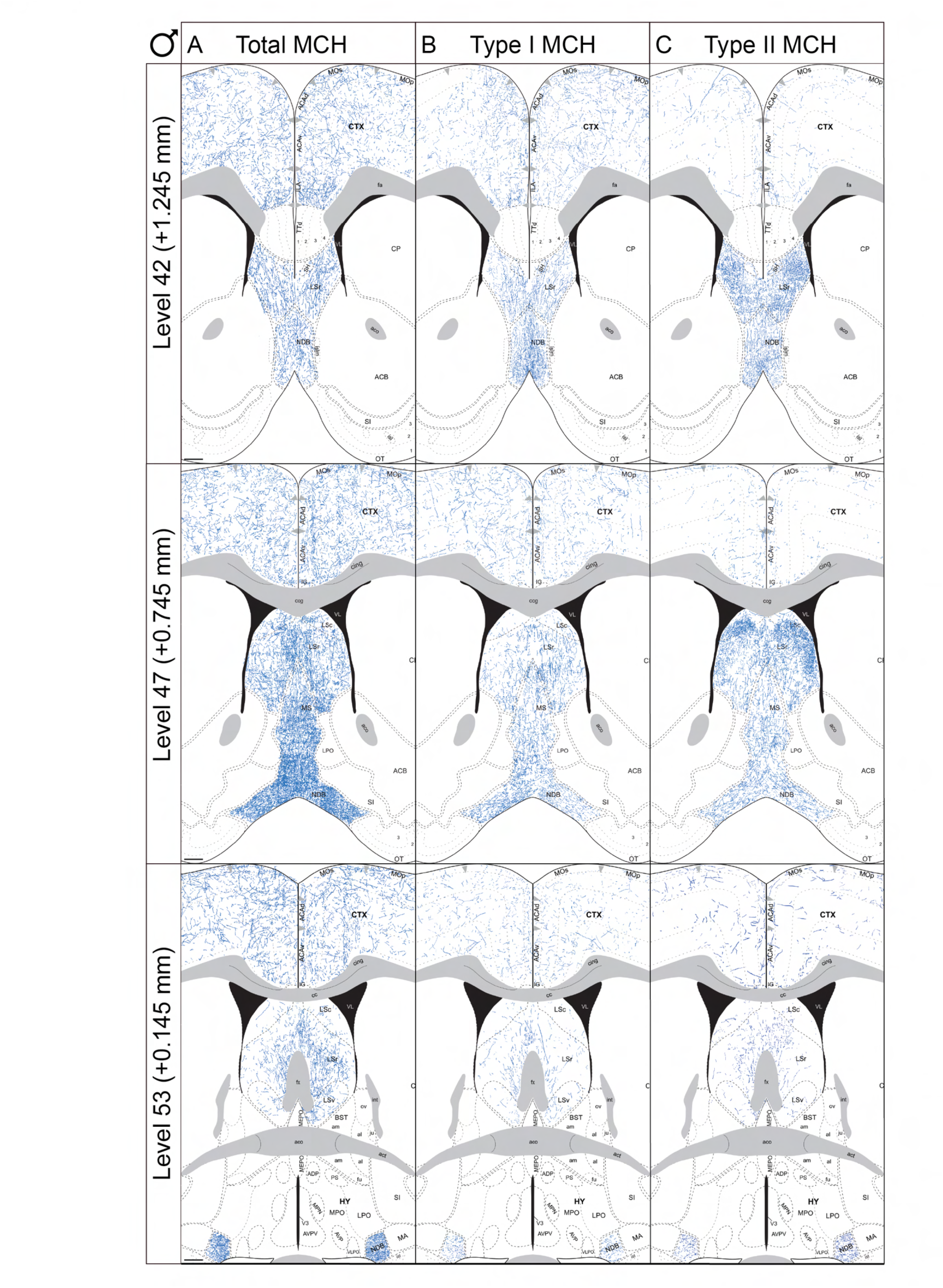
High resolution fiber mapping of total MCH, Type I MCH, and Type II MCH fibers in the septal complex and cortex in male mice. Representative coronal maps of GFP-immunoreactive fibers in representative anterior (top), middle (middle) and posterior (bottom) levels of the septal complex and adjacent cortex levels (distance from bregma: +1.245, +0.745, and +0.145 mm, respectively). Anatomical levels were identified by cytoarchitectural boundaries defined by the Allen Reference Atlas and only the MOs, MOp, ACA, LSr, LSc, LSv, MS, and NDB were traced.). Each column represents corresponding level traced from three experimental conditions: total MCH projections (**A**, *Pmch*-Cre+DIO-EYFP; left), Type I MCH projections (**B**, *Pmch*-Cre; *Tacr3*-Flpo +CV-Con/Fon-GFP; middle), and Type II MCH projections (**C**, *Pmch*-Cre; *Tacr3*-Flpo +CV-Con/Foff-GFP; right). **Abbreviations:** MOs: secondary motor area, MOp: primary motor area, ACAd: anterior cingulate area, dorsal part, ACAv: anterior cingulate area, ventral part, IG: indusium griseum, cing: cingulum bundle, CTX: cortex, CP: caudoputamen, ccg: corpus callosum (genu), LSc: lateral septal nucleus, caudal part, LSr: lateral septal nucleus, rostral part, LSv: lateral septal nucleus, ventral part, MS: medial septal nucleus, VL: lateral ventricle, LPO: lateral preoptic area, ACB: nucleus accumbens, SI: substantia innominata, NDB: nucleus of the diagonal band, OT: olfactory tubercle, BST: bed nuclei of the stria terminalis (ov: oval nucleus, al: anterolateral area, ju: juxtacapsular nucleus, am: anteromedial area), aco: anterior commissure (olfactory limb), MEPO: median preoptic nucleus, MPN: medial preoptic nucleus, ADP: anterodorsal preoptic nucleus, V3: third ventricle, AVPV: anteroventral periventricular nucleus, VLPO: ventrolateral preoptic area, MA: magnocellular nucleus, och: optic chiasm, isl: islands of Calleja, AVP: anteroventral preoptic nucleus, MPO: medial preoptic area, PS: parastrial nucleus, act: anterior commissure (temporal limb), fx: fornix. Scale bars represent 300 µM.

**Figure 18:**
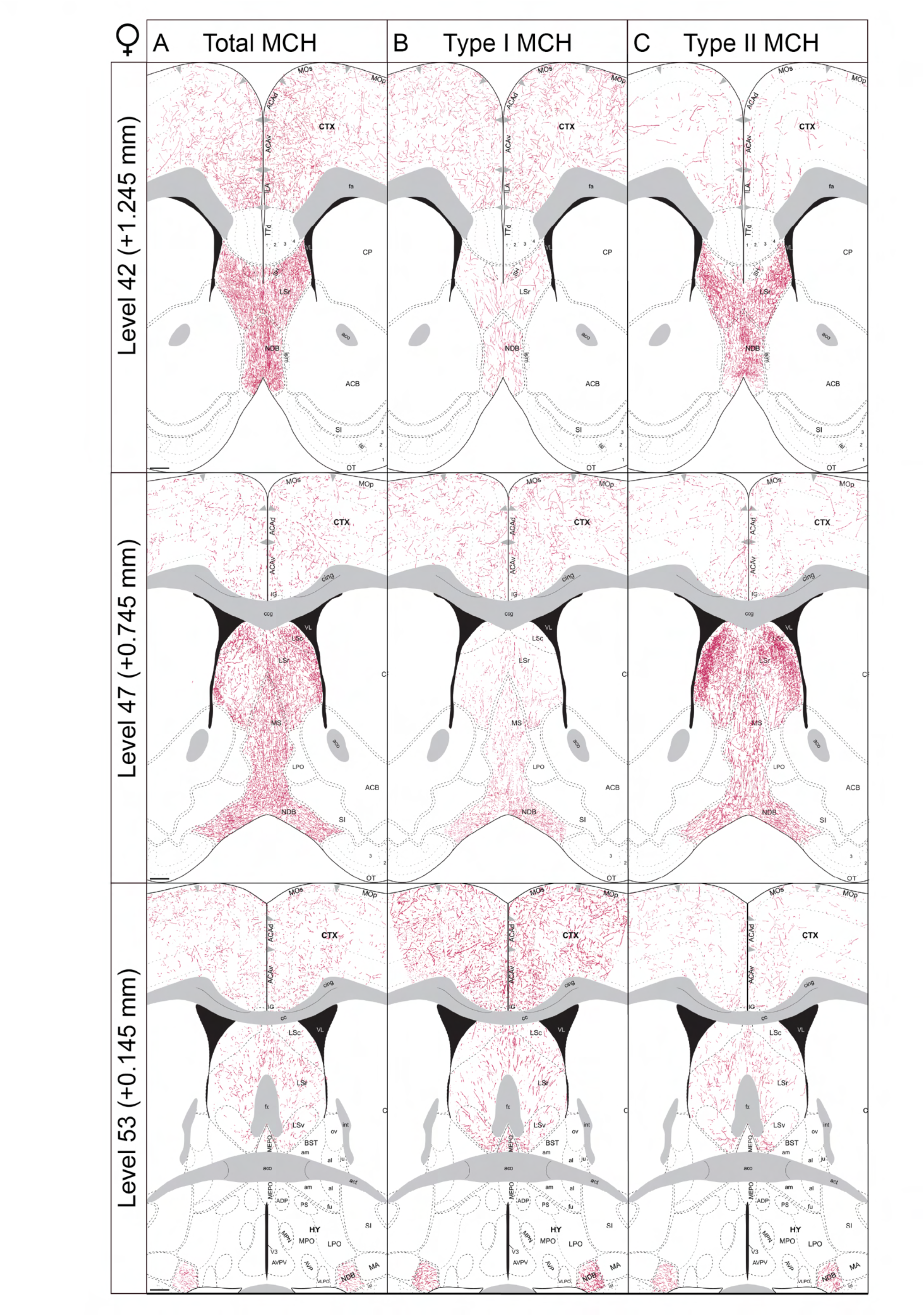
High resolution fiber mapping of total MCH, Type I MCH, and Type II MCH fibers in the septal complex and cortex in female mice. Representative coronal maps of GFP-immunoreactive fibers in representative anterior (top), middle (middle) and posterior (bottom) levels of the septal complex and adjacent cortex levels (distance from bregma: +1.245, 0.745, and +0.145 mm, respectively). Anatomical levels were identified by cytoarchitectural boundaries defined by the Allen Reference Atlas and only the MOs, MOp, ACA, LSr, LSc, LSv, MS, and NDB were traced.). Each column represents corresponding level traced from three experimental conditions: total MCH projections (**A**, *Pmch*-Cre+DIO-EYFP; left), Type I MCH projections (**B**, *Pmch*-Cre; *Tacr3*-Flpo +CV-Con/Fon-GFP; middle), and Type II MCH projections (**C**, *Pmch*-Cre; *Tacr3*-Flpo +CV-Con/Foff-GFP; right). **Abbreviations:** MOs: secondary motor area, MOp: primary motor area, ACAd: anterior cingulate area, dorsal part, ACAv: anterior cingulate area, ventral part, IG: indusium griseum, cing: cingulum bundle, CTX: cortex, CP: caudoputamen, ccg: corpus callosum (genu), LSc: lateral septal nucleus, caudal part, LSr: lateral septal nucleus, rostral part, LSv: lateral septal nucleus, ventral part, MS: medial septal nucleus, VL: lateral ventricle, LPO: lateral preoptic area, ACB: nucleus accumbens, SI: substantia innominata, NDB: nucleus of the diagonal band, OT: olfactory tubercle, BST: bed nuclei of the stria terminalis (ov: oval nucleus, al: anterolateral area, ju: juxtacapsular nucleus, am: anteromedial area), aco: anterior commissure (olfactory limb), MEPO: median preoptic nucleus, MPN: medial preoptic nucleus, ADP: anterodorsal preoptic nucleus, V3: third ventricle, AVPV: anteroventral periventricular nucleus, VLPO: ventrolateral preoptic area, MA: magnocellular nucleus, och: optic chiasm, isl: islands of Calleja, AVP: anteroventral preoptic nucleus, MPO: medial preoptic area, PS: parastrial nucleus, act: anterior commissure (temporal limb), fx: fornix. Scale bars represent 300 µM.

**Figure 19:**
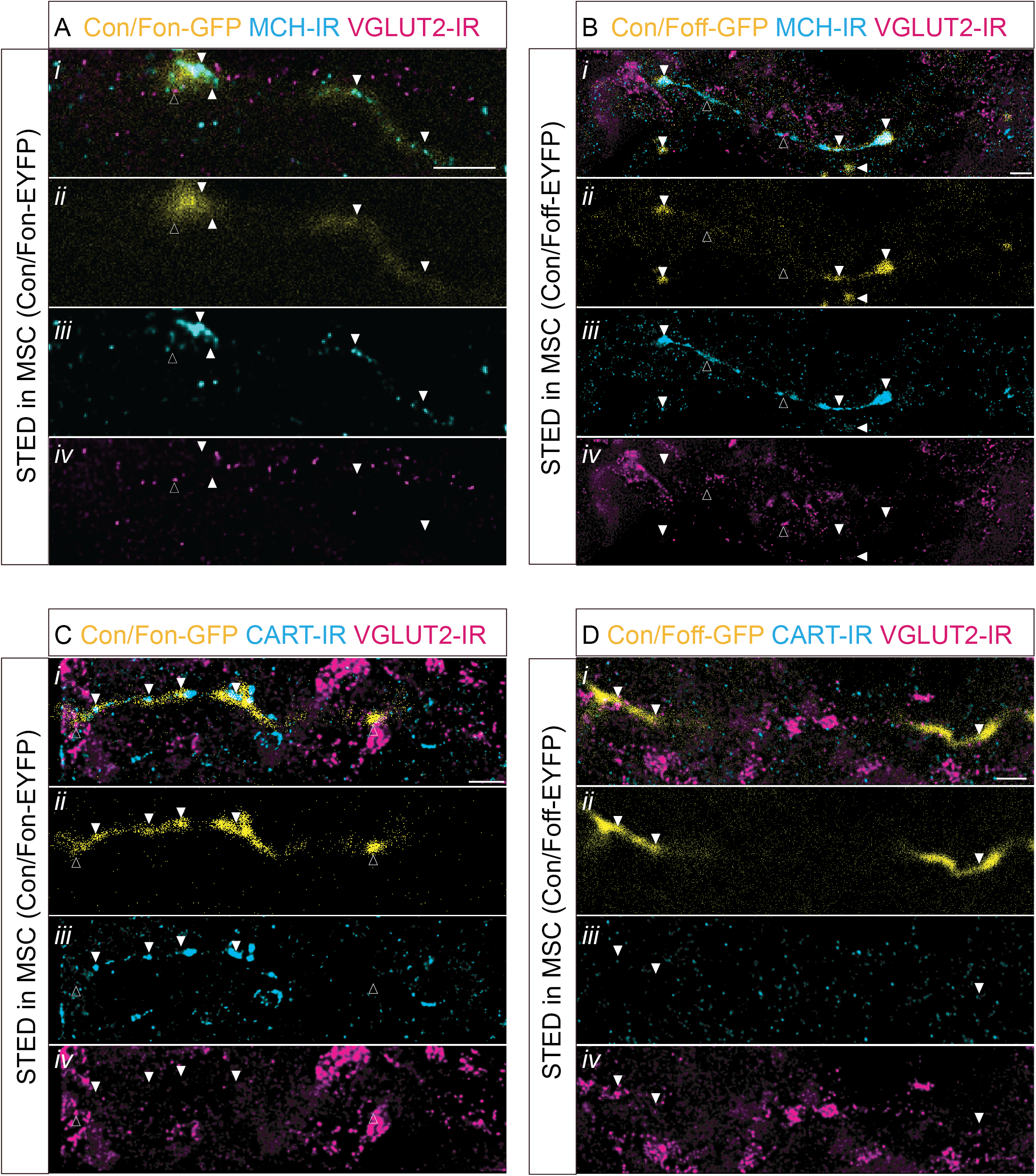
Super-resolution imaging reveals distinct neuropeptide profiles of CV-Con/Fon-GFP and CV-Con/Foff-GFP fibers axons in the medial septum and nucleus of the diagonal band. **A)** Representative merged channel (*i*) photomicrograph from CV-Con/Fon-GFP axons (yellow; *ii*) in the MSC showing MCH-IR (STED, cyan; *iii*) and VGLUT2-IR (magenta; *iv*) staining. **B)** Representative merged channel (*i*) photomicrograph showing CV-Con/Foff-GFP axons (yellow; *ii*) with MCH-IR (cyan; *iii*) and VGLUT2-IR (magenta; *iv*) staining in in the MSC. **C)** Representative merged channel (*i*) photomicrograph showing CV-Con/Fon-GFP axons (yellow; *ii*) with CART-IR (cyan; *iii*) and VGLUT2-IR (magenta; *iv*) staining in in the MSC. **D)** Representative merged channel (*i*) photomicrograph showing CV-Con/Foff-GFP axons (yellow; *ii*) with CART-IR (cyan; *iii*) and VGLUT2-IR (magenta; *iv*) staining in in the MSC. Panels *ii* (yellow) show confocal photomicrographs and panels *iii* (cyan) and *iv* (magenta) panels show stimulated emission depletion (STED) photomicrographs. White arrows indicate GFP colocalization with MCH-IR **(A, B)** or CART-IR **(C, D)**. Outlined arrows indicate GFP colocalization with VGLUT2-IR **(A-D)**. Scale bars represent 1 µM.

Overall, broad total MCH DIO-EYFP labeling was observed in the cortex, lateral septum (LS), medial septum (MS), and nucleus of the diagonal band (NDB) **(Figs. 17A, 18A)**. This recapitulates previous studies identifying strong MCH innervation of the cortex and LS (Bittencourt et al., 1992; Chee et al., 2015; Lima et al., 2013; Payant et al., 2024). Additionally, we did not observe any striking sex differences in fiber innervation, consistent with previous reports of comparable MCH subpopulation cell body count between male and female mice (Miller et al., 2024). Within midline cortical subregions, we observed broad fiber innervation throughout all layers of the primary motor area (MOp), secondary motor area (MOs), dorsal and ventral anterior cingulate area (ACAd,v) and infralimbic area (ILA). Fibers in cortical regions were best preserved in male and female CV-Con/Fon-GFP fibers in Type I MCH neurons and were only sparsely labeled by Con/Foff-EFYP virus in Type II MCH neurons.

Fiber innervation in the LS appeared to be denser in the caudal (LSc) and rostral (LSr) part of the LS, with fibers appearing to terminate or concentrate in the LSc. The LS was most densely innervated by Type II MCH neurons using the CV-Con/Foff-GFP virus, especially within the LSc. Both Type I and Type II MCH neurons exhibited fibers in the LSr but was more dense in the case of Type II MCH neurons. We found only sparse total MCH fibers in the ventral part of the LS (LSv).

In the MS and NDB, fibers from total MCH neurons were similarly dense throughout their anteroposterior extent in male or female mice. The MS and NDB exhibited equally dense innervation from both Type I and Type II MCH neurons.

In male and female CV-Con/Fon-GFP injected mice, ACA and MO fiber density appeared to be preserved, but a noticeable diminishment of fibers in the lateral septal complex when compared to total MCH fiber innervations **(Figs. 17B, 18B).** Interestingly, fibers in the medial septum and nucleus of the diagonal band also appeared to be preserved. Similar to the DIO-EYFP condition, Type I MCH fibers appeared sparser in ventral and posterior sections of the septal complex.

In contrast, CV-Con/Foff-GFP injected male and female mice showed dense septal innervation but a reduced fiber density in the cortex, when compares to the Type I MCH fiber distribution **(Figs. 17C, 18C).** Similar to the DIO-EYFP condition, lateral septal fibers appeared more dense in caudal and rostral subdivisions than ventral. Neither CV-Con/Fon-GFP nor CV-Con/Foff-GFP injected mice displayed obvious sex differences in fiber distribution or density within the septal and adjacent cortical subregions.

Together, these high spatial resolution maps demonstrate the complementary yet distinct projections of MCH subpopulations in septal and cortical areas. Type I MCH neurons preferentially innervate cortical structures with sparse septal innervation, while Type II MCH innervate the septal structures with sparse cortical innervation. However, both subsystems contribute similar innervations of the medial septum and nucleus of the diagonal band. The combined Type I and Type II fiber maps recapitulated the fiber pattern present in the DIO-EYFP condition that represented total MCH innervation. Overall, these distinct projection patterns were consistent between male and female mice, indicating that these subpopulation specific projection biases in the cortex and septum are likely conserved across sex.

### Super-resolution microscopy identifies molecular profiles of MCH subpopulations axon terminals in the MSC

Together, our comprehensive fiber anatomical mapping lays a foundation for understanding the functional output of MCH subpopulations, but the molecular identity of these axons remains poorly characterized. Although both Type I and Type II MCH neurons express MCH, only Type I MCH neurons are enriched in and likely release CART. MCH neurons have been shown to express *Vglut2* mRNA (Chee et al., 2015; Mickelsen et al., 2019) and protein (Del Cid-Pellitero & Jones, 2012; Qi et al., 2026), thus they have the capacity to co-release glutamate with MCH and/or CART along their axons. However, it is unknown if these molecular features are preserved at the axon terminal, and if MCH, CART, and VGLUT2 are differentially trafficked to Type I and Type II MCH varicosities. To address this, we performed Stimulated Emission Depletion (STED) super-resolution microscopy of MCH-IR, CART-IR, and VGLUT2-IR in CV-Con/Fon-GFP and CV-Con/Foff-GFP axons in the MSC, which is innervated by both populations of MCH neurons **(Fig. 19).**

We first surveyed MCH-IR and VGLUT2-IR at CV-Con/Fon-GFP and CV-Con/Foff-GFP axons **(Fig. 19A, B).** Qualitatively, while MCH-IR and VGLUT2-IR independently were observed within CV-Con/Fon-GFP and CV-Con/Foff-GFP axons, they rarely colocalized with one another **(Fig. 19A).** Additionally, we visualized sparse VGLUT2-IR in the vicinity of both CV-Con/Fon-GFP and CV-Con/Foff-GFP labeled axons.

We next observed CART-IR and VGLUT-IR at CV-Con/Fon-GFP and CV-Con/Foff-GFP axons **(Fig. 19C, D).** As expected, CART-IR was frequently found along CV-Con/Fon-GFP axons but not on CV-Con/Foff-GFP axons. Furthermore, VGLUT2-IR infrequently colocalized with CART-IR within both CV-Con/Fon-GFP and CV-Con/Foff-GFP axons.

Collectively, the super-resolution microscopy of the CV-Con/Fon-GFP and CV-Con/Foff-GFP axons in the MSC reveal that MCH subpopulation molecular identity is preserved in axons. Both Type I and Type II MCH axons contained MCH-IR, but CART immunoreactivity was more frequent in Type I axons. VGLUT2-IR was sparse along Type I and Type II MCH axons and rarely co-localized with MCH-IR or CART-IR, which suggests minimal co-packaging of glutamate with MCH or CART at the same release sites. Together with our anatomical mapping, these findings demonstrate that Type I and Type II MCH neurons differ not only in their projection patterns but also in the neurochemical composition of their axons, providing additional evidence for molecularly distinct MCH projections.

## DISCUSSION

In this study, we established and validated complementary intersectional genetic strategies to selectively access molecularly distinct MCH subpopulations and define their anatomical projection patterns. For intersectional targeting, we separated MCH neurons by *Tacr3* expression and operationally defined Type I (*Pmch*+/*Tacr3*+) and Type II (*Pmch*+/*Tacr3*-) MCH subpopulations. First, we generated the dual recombinase reporter *Pmch*-Cre;*Tacr3*-Flpo;RC::FLTG mouse to visualize Type I MCH cell bodies and axons and provide an initial framework for assessing subcircuits using our viral approach. However, this intersectional reporter strategy provided only moderate or sparse targeting of Type I MCH fibers. In our intersectional viral approach, we injected our dual recombinase *Pmch*-Cre;*Tacr3*-Flpo line with ConVERGD viruses; using the CV-Con/Fon-GFP virus to label Type I MCH neurons and the CV-Con/Foff-GFP virus to label Type II MCH neurons. Furthermore, we also used ConVERGD viruses in parallel with a DIO-EYFP virus in *Pmch*-Cre mice to compare axonal projection patterns of Type I and II MCH neurons with those of all MCH neurons.

Here, we identified that Type I and Type II MCH neurons extend fibers to overlapping brain regions, but they each also exhibit preferential axonal projection patterns. Type I MCH neurons preferentially project towards cortical structures, accounting for much of the total MCH innervation of the cortex, as well as projecting to many subcortical structures. Type II MCH neurons, in turn, exhibit very sparse innervation of the cortex with preferential innervation of certain subcortical structures, including the lateral septum and inferior colliculus among others. Our high-resolution fiber maps further revealed that the distribution of fibers within these regions were not uniform and exhibited distinct spatial distribution patterns, for example within the LSc versus LSr. Finally, using super-resolution microscopy, we identified MCH-IR, CART-IR, and/or VGLUT2-IR in Type I and Type II MCH axons, recapitulating the molecular profile of these subpopulations.

### Development and validation of intersectional genetic tools for MCH subpopulation access

Previous studies have been limited in their ability to access and manipulate spatially intermixed Type I and Type II MCH neuronal subpopulations. Here, we established three complementary tools for the manipulation of MCH subpopulation-specific investigation: the novel *Tacr3*-Flpo line for genetic identification of Type I MCH neurons, intersectional anterograde ConVERGD viruses for subcircuit anatomical tracing, and the dual recombinase reporter *Pmch*-Cre;*Tacr3*-Flpo;RC::FLTG line for independent validation of Type I MCH projection patterns.

We also validated Conditional Viral Expression by Ribozyme Guided Degradation (ConVERGD) viruses for the targeting of MCH subpopulations in our dual recombinase *Pmch*-Cre;*Tacr3*-Flpo model using control injections and immunohistochemistry. Through this approach, we identified high specificity and penetrance of CV-Con/Fon-GFP virus for targeting Type I MCH neurons and CV-Con/Foff-GFP virus for targeting Type II MCH neurons. Compared to the well-described INTRSECT (INTronic Recombinase Sites Enabling Combinatorial Targeting) viruses (Fenno et al., 2014, 2020), ConVERGD viruses (Hughes et al., 2024) offer an alternative viral approach that uses a smaller and simpler viral design and self-cleaving riboyzmes, rather than recombinase-based intron engineering, to prevent non-specific expression. Anecdotally, in our *Pmch*-Cre;*Tacr3*-Flpo model, ConVERGD viruses provided stronger specificity and a higher recombination payload than INTRSECT viruses (data not shown).

Additionally, our results provide the framework for the application of ConVERGD viruses in interrogating MCH subcircuit function. Beyond the GFP construct we validated, ConVERGD viruses can be packaged with Designer Receptors Exclusively Activated by Designer Drugs (DREADDs), light-gated opsins, or genetically encoded calcium indicators (GCaMPs) (Hughes et al., 2024). As a result, these tools would enable monitoring of MCH subpopulation activity across behavioral states or selective activation or inhibition of subcircuits during behavioral paradigms.

Finally, we validated the specificity and penetrance of the triple mutant *Pmch*-Cre;*Tacr3*-Flpo;RC::FLTG to complement our viral strategy. Upon successful dual recombination by Cre and Flpo, this line will drive eGFP expression in Type I MCH neurons (Plummer et al., 2015). While we identified that this line reported fewer Type I MCH neurons (42.3 ± 3.6%) than in our *Tacr3*-Flpo;tdT mouse (60.3 ± 2.7%), it exhibited high specificity for Type I MCH neurons (98.0 ± 0.6%). Its reduced penetrance likely reflects the requirement of two independent recombination events to occur within the same cell, with each requiring sufficient levels of recombinase expression to drive eGFP fluorescence. If either Flpo or Cre recombinase is expressed at insufficient levels when the target locus is available (i.e., temporal mismatching in the development of Cre and Flpo expression), the stop cassette will prevent eGFP fluorescence (Plummer et al., 2015). A practical limitation of this approach is that this line serves solely as a reporter for Type I MCH neurons, which restricts its utility to solely visualize these neurons without functional manipulation.

Despite these limitations, our validated intersectional viral and genetic approaches in accessing MCH subpopulations provides robust, cross-validated identification of Type I and Type II MCH neurons and establishes the necessary foundation for future work in investigating subcircuit function.

### Functional implications of molecularly defined MCH subcircuits

MCH neurons have been implicated in a broad range of physiological and behavioral processes, including sleep-wake regulation, feeding, memory, reward, and motivated behaviors (Adamantidis & de Lecea, 2009; Al-Massadi et al., 2021; Bandaru et al., 2020; Concetti et al., 2023; Diniz & Bittencourt, 2017; Lord et al., 2021). Recent evidence supports functional heterogeneity within the MCH system, with miniscope and fiber photometry recordings of MCH neurons identifying distinct wake- and sleep-active MCH subpopulations (Blanco-Centurion et al., 2019; Izawa et al., 2025). Additionally, distinct subsets of MCH neurons respond differently to afferent signals such as insulin (Hausen et al., 2016) and hypocretin/orexin (Izawa et al., 2025). Finally, recent work also indicated that MCH neurons may participate in functionally distinct subcircuits, as activation of nucleus accumbens MCH projections enhance the reward value of food, but has no effect on baseline feeding or REM sleep (Furman et al., 2025). Although previous studies have established fundamental roles for MCH neurons in diverse physiological and behavioral processes, the absence of approaches for selectively accessing genetically defined MCH subpopulations has limited investigation of their potentially distinct circuit functions.

Using complementary intersectional genetic strategies, we demonstrate that molecularly defined Type I and Type II MCH neurons comprise largely overlapping, yet preferentially organized, projection networks. While both subpopulations innervate many common targets, Type I MCH neurons preferentially project to cortical and hippocampal regions, including the orbital area, anterior olfactory nucleus, anterior cingulate area, primary motor area, primary somatosensory areas, piriform area, and dorsal and ventral hippocampal formation. Several of these cortical and hippocampal subregions have been attributed to roles in locomotion, memory consolidation, sleep, olfaction, and impulsivity which have been associated with the MCH system as whole (Adams et al., 2011; Hausen et al., 2016; Izawa et al., 2019; Monzon et al., 1999; Noble et al., 2019; Pissios et al., 2008).

Type I MCH neurons robustly express the neuropeptide CART (Broberger, 1999; Elias et al., 2001; Fujita et al., 2021; Izawa et al., 2025; Mickelsen et al., 2019; Miller et al., 2024) and likely have the capacity to release CART as a co-modulator at synaptic sites (Jancsik et al., 2018). LHA CART has been shown to be orexigenic, as reintroduction of CART expression in LHA CART deficient mice increases food consumption and fat mass (Lau et al., 2017). However, we predict that Type I MCH neurons exert functionally distinct effects depending on the target region. For example, site-specific delivery of CART can produce anorectic effects via the lateral septum. Infusion of MCH into the lateral septum increases feeding, and co-infusion of CART suppresses MCH-mediated feeding at the LS (Shankhatheertha et al., 2025). While Type I MCH neurons exhibited denser fiber tracts in the medial aspect of the LSr, the anorexigenic effect of CART was more prominent within the lateral aspect of the LSr (Shankhatheertha et al., 2025) thus suggesting that CART may diffuse to distal but neighboring sites of action.

Additionally, Type I MCH neurons also express functional NK3R (encoded by *Tacr3* gene) and depolarize in response to a neurokinin-B (NKB) analog (Fujita et al., 2021). We previously identified that tachykinergic neurons stemming from the central extended amygdala provide broad GABAergic input onto MCH neurons (Fujita et al., 2021). As these neurons also produce NKB, this pathway may preferentially influence Type I MCH neurons. Future studies will be required to determine whether this molecular specificity contributes to functional differences between MCH subpopulations.

Type II MCH neurons preferentially innervate septal, thalamic, midbrain, and hindbrain structures, with moderate to dense fibers present in the septal complex, medial preoptic area, nucleus reuniens, mammillary body, periaqueductal grey, ventral tegmental area and superior colliculus. Previous manipulations of MCH projections in these regions resulted in disruptions in REM sleep (Jego et al., 2013; Kroeger et al., 2019) and memory processing (Liu et al., 2022). It is also notable that Type II MCH neurons also preferentially innervate the lateral septum, which has been implicated in the regulation of a wide range of other behaviors including navigation, locomotion (Wirtshafter and Wilson, 2019), feeding and other motivated behaviors (Sweeney and Yang, 2016; Rizzi-Wise and Wang, 2021; Dib et al, 2025; Payant et al, 2024; Goode et al., 2026).

Together, the projection patterns of Type I and Type II MCH recapitulated the overall pattern of MCH fiber distribution described in previous anatomical studies (Bittencourt et al., 1992) and observed in our DIO-EYFP mapping of total MCH projections. Rather than supporting completely segregated anatomical circuits, these findings suggest that MCH subpopulations are organized as partially overlapping networks with distinct regional biases, providing a potential anatomical substrate through which genetically defined MCH neurons may differentially influence downstream targets. However, whether these subcircuit projection biases may translate into subpopulation-specific functional contributions remains to be determined.

### Type I and Type II MCH axons exhibit distinct neuropeptides signatures in the MSC

Using super-resolution microscopy, we observed MCH-IR, CART-IR, and VGLUT2-IR in CV-Con/Fon-GFP and CV-Con/Foff-GFP axons in the MSC. Type I CV-Con/Fon-GFP fibers typically contained MCH and CART, while Type II CV-Con/Foff-GFP fibers typically contained MCH but little CART. This underlies the molecular phenotype of Type I MCH neurons that are enriched in CART. Together, these data suggest that Type I and Type II MCH neurons may exert differential neuromodulatory effects through the release of MCH or CART.

Importantly, our work further confirms the glutamatergic identity of MCH neurons. While MCH neurons express VGLUT2 (Abrahamson et al., 2001; Harthoorn et al., 2005; Mickelsen et al., 2017, 2019) and have been shown to release glutamate in the LS (Chee et al., 2015; Liu et al., 2022), they also express GAD65/67 (Elias et al., 2001; Harthoorn et al., 2005; Sapin et al., 2010) and GABA release has been observed following optogenetic activation of MCH fibers in the tuberomammillary nucleus (Jego et al., 2013). Additionally, a very small portion of fibers from MCH neurons in the LC co-localize with VGAT, the vesicular transporter for GABA (Del Cid-Pellitero & Jones, 2012). As a result, the fast neurotransmitter profile of MCH neurons has been debated. Our study did not assess GABAergic markers, so the capacity of MCH neurons to mediate GABAergic signaling remains an open question. Here, we provide another line of evidence that MCH neurons likely participate in glutamatergic signaling within the MSC. We found sparse VGLUT2-IR colocalization along MCH-IR axons that is consistent with previous functional work (Chee et al., 2015; Liu et al., 2022) and similar to the observation of low density VGLUT2-IR localization in MCH axons in the LC (Del Cid-Pellitero & Jones, 2012) and olfactory bulb (Qi et al., 2026). Furthermore, VGLUT2 expression at axon terminals may not be readily labeled by virally-transduced reporter expression, thus VGLUT2 and MCH colocalization would be under-reported.

### Additional limitations and future directions

A central finding emerging from our neuroanatomical analyses is that Type I and Type II MCH neurons form overlapping rather than anatomically discrete or mutually exclusive subcircuits. Additionally, many subregions have equivalent converging projections from Type I and Type II MCH neurons, for example in the nucleus accumbens (ACB), lateral mammillary nucleus (LM), superior colliculus (SC), amygdala, and several brainstem nuclei. Whether Type I and Type II MCH neurons exert synergistic or antagonistic effects in these subregions of converging input is not yet known, thus limiting our speculation on the function of each MCH subcircuit. Future studies combining ConVERGD-based targeting with genetically encoded calcium indicators, fiber photometry, and other functional approaches could determine how the overlapping projection networks of Type I and Type II MCH neurons influence downstream circuits and are differentially recruited across diverse physiological and behavioral states. Such studies could help explain how molecular and circuit heterogeneity within the MCH system may contribute to its diverse roles in REM sleep, feeding, exploration, and motivated behavior.

Second, our model relies on the binary separation of MCH subpopulations based on the expression of *Tacr3*, which necessarily simplifies the underlying transcriptional heterogeneity of these subpopulations. Our single-cell RNA sequencing identified *Tacr3*, together with *Cartpt*, as significantly enriched in one transcriptionally defined MCH subpopulation, although expression of both genes extends across the transcriptional boundary between populations (Mickelsen et al., 2019). Thus, *Tacr3* and *Cartpt* may be better considered markers of differential enrichment along a continuum of gene expression rather than categorical markers of transcriptionally or functionally discrete MCH subpopulations.

Third, while minimal, leakiness in viral expression was reported. While ConVERGD viral specificity was high, there was still a proportion of CV-Con/Fon-GFP and CV-Con/Foff-GFP that did not express MCH-IR (9.1% and 10.5%, respectively). While this is unlikely to alter overall projection patterns that we observed, it should be considered when interpreting fiber labeling in regions with sparse fiber presence. Additionally, 26.3% of CV-Con/Foff-GFP co-expressed CART-IR, indicating possible non-specific expression in Type I MCH neurons. This may conflate a small proportion of Type I and II fiber innervation.

Fourth, while our intersectional viral and genetic reporter strategies define the anatomical distribution of MCH subpopulation projections, they do not distinguish functional synaptic terminals from fibers of passage. Future studies combining these intersectional tools with functional circuit-mapping approaches will be important for identifying functional synaptic connections at downstream targets. In parallel, the use of synaptically-targeted fluorescent reporters may facilitate visualization of subpopulation-specific presynaptic terminals and further refine the anatomical organization of these projections.

Overall, despite these limitations, our complementary intersectional genetic and viral approaches establish the first subpopulation-resolved anatomical framework for genetically defined MCH neurons. By revealing that Type I and Type II MCH neurons comprise largely overlapping yet preferentially organized projection networks, this work provides a foundation for future studies investigating how molecularly defined MCH subpopulations differentially contribute to neural circuit function and behavior.

## ACKNOWLEDGEMENTS

We gratefully acknowledge the members of the Jackson, Chee, and Schwarz labs for support, assistance and helpful discussions, C. Kangas for genotyping support, and C. O’Connell and J. Martin for imaging support. Many thanks to the NIMH Transgenic Core for generating the *Tacr3*-Flpo mouse line. We are grateful to P. Jensen and N. Sciolino for generously sharing the RC::FLTG mouse line. Project supported by NIH grants R01MH112739 and R01NS131664 (to ACJ), NSERC Discovery Grant RGPIN-2024-04133 (to MJC), institutional funds from St. Jude Children’s Research Hospital (to LAS), and an NIH Shared Instrumentation Grant S10OD023618 (to C. O’Connell) and S10OD016435 to (A. Nishiyama).

---

**Table 1:** Table of Primary Antibodies.

| **Name** | **Vendor** | **Catalog No.** | **RRID** |
| --- | --- | --- | --- |
| Rabbit anti-MCH | Phoenix Pharmaceuticals | H-070-47 | AB_2722682 |
| Rabbit anti-CART | Phoenix Pharmaceuticals | H-003-61 | AB_2922768 |
| Chicken anti-GFP | Invitrogen | A10262 | AB_2534023 |
| Rabbit anti-NK3R | Novus Biologicals | NB300 | AB_10003270 |
| Chicken anti-VGLUT2 | Synaptic Systems | 135 409 | AB_2943526 |

**Table 2:** Table of Secondary Antibodies.

| **Name** | **Vendor** | **Catalog No.** | **RRID** |
| --- | --- | --- | --- |
| Donkey anti-Rabbit Alexa Fluor 488 | Abcam | ab150073 | AB_2636877 |
| Donkey anti-Rabbit Alexa Fluor 594 | Abcam | ab150076 | AB_2782993 |
| Donkey anti-Rabbit Alexa Fluor 647 | Abcam | ab150083 | AB_2714032 |
| Goat anti-Chicken Alexa Fluor 488 | Abcam | ab150169 | AB_2636803 |
| Goat anti-Rabbit STAR ORANGE | Aberrior | STORANGE-1002-500UG | AB_3068622 |
| Goat anti-Chicken STAR RED | Aberrior | STRED-1005 | N/A |

**Table 3:** Table of RNAscope Probes.

| **Name** | **Vendor** | **Catalog No.** |
| --- | --- | --- |
| RNAscope™ Probe-Mm-Tacr3 | Advanced Cell Diagnostics | 481671 |
| RNAscope™ Probe-pFlpO | Advanced Cell Diagnostics | 520791 |

